# Global transcriptomic profiling of the bone marrow stromal microenvironment during postnatal development, aging and inflammation

**DOI:** 10.1101/730457

**Authors:** Patrick M. Helbling, Elena Piñeiro-Yáñez, Rahel Gerosa, Steffen Boettcher, Fátima Al-Shahrour, Markus G. Manz, César Nombela-Arrieta

## Abstract

Bone marrow (BM) stromal cells provide the structural and regulatory framework for hematopoiesis and contribute to developmental-stage specific niches, such as those preserving hematopoietic stem cell (HSCs). Despite recent advances in our understanding of stromal composition and function, little is known on the dynamic transcriptional remodeling that this compartment undergoes over time and during adaptation to stress. Similarly, how molecular changes in stroma are linked to age-related modulation of hematopoiesis is poorly understood. Using RNA-sequencing, we performed a longitudinal comparison of the transcriptional profile of four principal mesenchymal and endothelial stromal subsets, namely CXCL12-abundant reticular (CARc), PDGFR-α^+^Sca-1^+^, sinusoidal (SECs) and arterial endothelial cells (AECs), isolated from early postnatal, adult and aged mice. Our data i) provide molecular fingerprints defining novel, cell-specific anatomical and functional features ii) reveal radical reprogramming of pro-hematopoietic, immune and matrisomic transcriptional programs during the narrow temporal transition from juvenile to adult stages iii) demonstrate that homeostatic aging is characterized by a progressive and pronounced upregulation of pro-inflammatory gene-expression and loss of stromal cell fitness. By profiling *in vivo* responses of stromal cells to infection-mimicking agents, we finally demonstrate that transcriptomic pathways elicited by sterile inflammation are largely recapitulated during aging, thereby supporting the inflammatory basis of aging-related adaptations of BM hematopoietic function.

## Introduction

The continuous production of massive amounts of blood cell lineages is an absolutely essential process, which is centralized in bone marrow (BM) tissues during adulthood and is maintained by hematopoietic stem cells (HSCs) (Eaves, 2015; Nombela-Arrieta and Manz, 2017). Throughout life, BM function quantitatively and qualitatively adjusts to match the fluctuating demands of organisms (Takizawa et al., 2012). For instance, during early postnatal development, HSC populations substantially expand through self-renewing proliferation to cope with the rapid growth in blood and marrow volume, and the expansion of the hematopoietic system characteristic of these phases. With acquisition of definitive body dimensions, growth is slowed down and HSCs sharply transition from active self-renewing proliferation to a hibernating state, through a developmentally-timed switch, which in mice happens at 3-4 weeks of age (Benz et al., 2012; Bowie et al., 2006). In absence of overt disease or insults, adult HSCs will thereafter remain largely quiescent and keep only sporadic cycling activity, sufficient to support the homeostatic maintenance of the HSC pool and hematopoietic production (Bernitz et al., 2016; Nakamura-Ishizu et al., 2014; Takizawa et al., 2011; Wilson et al., 2008). Beyond marked differences in proliferative rates, juvenile and adult HSCs differ in transcriptional regulation and lineage potential (Kim et al., 2007; Ye et al., 2013). Furthermore, compared to adult phases, neonatal BM tissues display important differences in erythropoietic production and mature B cell content (Moscatello et al., 1998; Pihlgren et al., 2001).

Physiological aging also encompasses various alterations in hematopoiesis and immune cell function. Aging leads to a gradual decrease in lympho- and erythropoiesis and a concomitant bias towards myeloid cell production (Geiger et al., 2013). In aged mice, HSCs significantly expand but exhibit deficits in repopulating capacity, homing ability, aberrant lineage differentiation and activation of intracellular stress pathways (Beerman et al., 2010; de Haan and Lazare, 2018). Several aging-related traits seem to recapitulate, albeit with lower magnitude, those caused by pathogenic or inflammatory challenges (Kovtonyuk et al., 2016). As an example, bacterial infections elicit expansion of progenitor cell subsets, biased and enhanced production of granulocytes (emergency granulopoiesis), and mobilization of HSCs to extramedullary sites (Manz and Boettcher, 2014). Viral agents also trigger BM hypoplasia, drive monopoiesis and impair erythropoiesis and B lymphopoiesis (Nombela Arrieta and Isringhausen, 2017). A common hallmark of hematopoietic responses to inflammatory challenges is the rapid activation of HSC cycling, which increases their proliferative history and leads to functional deficits (Takizawa et al., 2017; Walter et al., 2015; Zhang et al., 2016). The existing similarities between aging and inflammation-related effects have inspired the *inflammaging* hypothesis, which associates age-induced degeneration to the progressive emergence of a low-grade, basal inflammatory condition at tissue wide and systemic levels (Kovtonyuk et al., 2016).

The non-hematopoietic BM stromal compartment modulates hematopoiesis through direct crosstalk and local delivery of regulatory signals (Mercier et al., 2012; Nombela-Arrieta and Manz, 2017). A number of mesenchymal cell subsets have been described, with proven functional roles in the orchestration of homeostatic and reactive hematopoiesis (Kfoury and Scadden, 2015). Thus far, the most extensively characterized stromal subset is formed by so-called CXCL12 abundant reticular cells (CARc), which are ubiquitous in adult BM, overlap to a high degree with Leptin receptor-expressing (LepR^+^) cells and can be phenotypically defined as CD45^−^Ter119^−^CD140b^+^Sca-1^−^ cells. CARc express high levels of crucial soluble regulators such as stem cell factor (SCF), pleiotrophin (PTN) or interleukin-7 (IL-7) (Ding et al., 2012; A. C. Gomes and M. S. Gomes, 2016; Himburg et al., 2018) and have been assigned many roles in the direct regulation of HSCs, lympho- and myelopoiesis (Ding et al., 2012; Greenbaum et al., 2013; Kohara et al., 2007; Noda et al., 2011; Omatsu et al., 2010; Sugiyama et al., 2006; Tokoyoda et al., 2004). Originally considered as a single cellular entity on the basis of the high expression of GFP in *Cxcl12*-GFP mice (Ara et al., 2003), recent single cell studies clearly show that CARc are heterogeneous and comprised of at least 4 subpopulations, which form a differentiation continuum of osteo- and adipogenic progenitors (Baryawno et al., 2019; Tikhonova et al., 2019). Additional BM mesenchymal stromal subsets have been characterized with less detail (Kfoury and Scadden, 2015). Among them, PDGFR-α^+^Sca-1^+^ (here designated PαSc) were initially described on the basis of a phenotypic signature that uniquely differs from that of CARc on the expression of Sca-1 (Morikawa et al., 2009). Flow cytometry studies have shown that PαS cells are found at highest frequencies in developing bone, but gradually decline to represent a very low fraction of the mesenchymal compartment in adult BM (Hu et al., 2016; Nusspaumer et al., 2017). Both CARc and PαSc contain CFU-F potential, trilineage differentiation capabilities *in vitro*, and give rise to osteoblasts, reticular stroma and adipocytes *in vivo* (Morikawa et al., 2009; Omatsu et al., 2010). However, the ontogenetic relationships and functional roles in skeletal development and repair of both populations are unclear. Furthermore, although some studies have suggested a potential contribution of PαSc to the regulation of HSCs (Greenbaum et al., 2013; Hu et al., 2016), their anatomical distribution and regulatory roles in hematopoiesis remain largely unexplored.

Endothelial cells are also an integral part of the BM microenvironment. To date BM microvessels have been divided in three main vascular districts based on structural, topological and phenotypic criteria (Bixel et al., 2017; Sivaraj and Adams, 2016). Arterial circulation, made up of large, centrally running arteries and arterioles of smaller caliber, gives rise to sinusoidal circulation through so-called transitional vessels (TV) (Kusumbe et al., 2016). Arterial and sinusoidal endothelial cells (AECs and SECs) can be distinguished by specific marker expression (Gomariz et al., 2018; X. M. Li et al., 2009; Smith-Berdan et al., 2015), express varying levels of SCF and CXCL12 and regulate different aspects of hematopoiesis (Ding et al., 2012; Xu et al., 2018), from HSC maintenance to maturation of megakaryocytes and late B cell differentiation (Lazzari and Butler, 2018; Stegner et al., 2017).

Although major advances have been made in the understanding of the composition and structure of the stromal compartment in homeostasis, much less is known about the molecular changes that stromal cells undergo during the aging process over time and during hematopoietic challenges, and how these may be potentially linked to adaptations of hematopoietic function. Here we present a longitudinal analysis of the evolution of genome-wide expression profiles of CARc, PαSc, AECs and SECs across three representative stages of postnatal lifespan and during inflammatory responses to prototypical infection-mimicking agents.

## Results

### Stromal cell isolation, transcriptional profiling and validation

We employed previously validated phenotypic signatures to prospectively isolate pure cellular populations of CARc, PαSc, SECs and AECs using fluorescence activated cell sorting (FACS) (Figure 1A, 1B). Within the endothelial cell population (CD45^−^Ter119^−^CD31^hi^), SECs and AECs were identified as Sca-1^int^CD105^hi^ and Sca-1^hi^CD105^int^, respectively (Figure 1B) (Gomariz et al., 2018). Although TVECs have been recently described (Kusumbe et al., 2014), a molecular signature to reliably distinguish these cells has not been found even using sc-RNA-Seq analyses (Baryawno et al., 2019; Tikhonova et al., 2019). Nonetheless, based on immunohistological data, TVECs are relatively rare, they express low/intermediate levels of CD105, and therefore are most likely isolated as AECs in our gating strategy (Gomariz et al., 2018; Itkin et al., 2016; Kusumbe et al., 2014). In turn, mesenchymal CARc and PαSc were sorted from the non-hematopoietic (CD45^−^Ter119^−^), non-endothelial (CD31^−^) compartment as CD140b^+^Sca-1^−^ (CARc) and CD140b^+^Sca-1^+^ (PαSc) (Figure 1B) (Morikawa et al., 2009). To get a global overview of age-dependent transcriptional oscillations in BM stroma, we isolated all four subsets from 2 week (2w - juvenile), 2 months (2m - adult) and 2 years (2y - aged) old mice. All populations were consistently present and their phenotypic signatures largely conserved at different ages (Figure S1A). We sorted 4000-50’000 cells for each cell type and condition and performed high-throughput RNA-seq in at least three independent biological replicates per experimental condition. Transcripts corresponding to non-coding RNAs, pseudogenes and genes below the detection threshold were filtered out, resulting in the identification of a total of 11’595 protein coding genes in all conditions (full list provided in Table S1).

**Figure 1.**
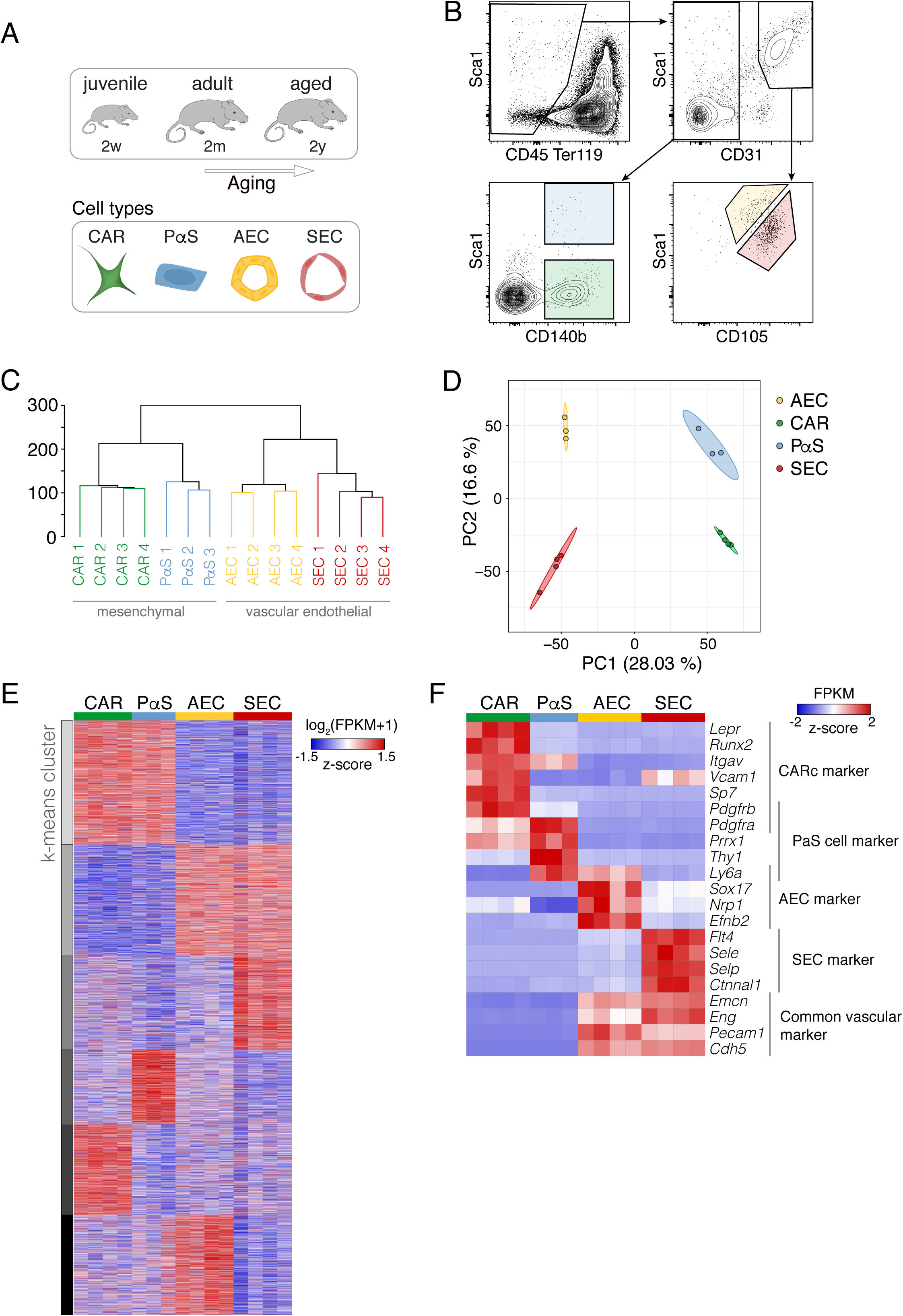
Global transcriptomic analysis of juvenile, adult and aged BM stroma. **A.** Schematic graphical overview displaying the four stromal cell types isolated from mice of three different ages that were investigated in this study. CAR: Cxcl12-abundant reticular cells. PαS: CD140b^+^ Sca1^+^ mesenchymal cells. AEC: arterial endothelial cells. SEC: sinusoidal endothelial cells. **B.** Fluorescence activated cell sorting (FACS) gating strategy for BM stromal cell populations (color-coded) in the adult condition. Scatterplots depict cells pre-gated for singlets and live cells (DAPI^−^). **C and D.** Unsupervised hierarchical clustering (dendrogram) and principal component analysis (PCA) of the four populations in the adult condition based on the expression values (FPKM) of all detected genes. Numbers 1-4 in (**C**) and individual dots in (**D**) represent independent biological replicates. **E.** K-means clustering analysis using all detected genes in stromal cells of 2m old mice, to separate genes into 6 clusters. **F.** Heatmap of expression previously reported stromal cell type-specific markers. Row z-score normalized expression values.

Unsupervised hierarchical clustering and principal component (PCA) analyses revealed very high reproducibility between sample replicates (Figure 1C, 1D, 1E and S1B). Consistent with their shared cellular identity and developmental origins, endothelial and mesenchymal components separated most from each other, while AECs and SECs, as well as CARc and PαSc respectively clustered together (Figure 1D). Inspection of expression values of cell-type specific known markers, including those employed for their identification and cell sorting, confirmed the robustness of the isolation strategies and cell type identities (Figure 1F). As expected, CARc expressed high levels of *Cxcl12*, *Lepr*, *Itgva*, *Vcam-1*. Genes associated to mesenchymal identity such as *Pdgfra*, *Pdgfrb*, and *Prrx1* were commonly expressed by CARc and PαSc. Transcripts of prototypical vascular endothelial proteins (*Pecam1*, *Tek* and *Cdh5*) were exclusively and abundantly detected in SECs and AECs. Alternative markers previously described to discriminate sinusoidal from arterial vessels in immunohistological studies also displayed highly specific expression of SECs or AECs in our dataset. Arterial-specific *Sox17* and *Ephb2* were exclusively and strongly detected in AECs, while *Selp* (P-Selectin), *Sele* (E-Selectin), *Ctnnal1* (Alpha-catulin) and *Flt4* (VEGFR-3) were present in the SECs fractions (Acar et al., 2015; Butler et al., 2010; Corada et al., 1AD; Gomariz et al., 2018). Notably, the specificity in expression of the vast majority of markers was maintained at all timepoints (Figure S1C). Furthermore, samples from each population consistently clustered throughout developmental stages, indicating preservation of overall cellular identity across lifespan (Figure S1B).

### Cell-subset specific transcriptomic fingerprints for phenotypic detection

We first mined RNA-seq data from BM from 2-month old mice to uncover potential novel genes selectively expressed by a particular cell type in homeostasis. To this end we established restrictive filtering criteria and focused on genes displaying i) low/negligible signal (FPKM values < 30) in three stromal cell types ii) 10-fold increase (log_2_-fold change (FC) > 3.32, FDR ≤ 0.05) in the remaining cell type with respect to all other subsets. This yielded transcriptomic fingerprints containing 55, 55, 24 and 38 highly specific genes for CARc, PαSc, AECs and SECs, respectively (Figure S2A). The CARc signature was dominated by genes associated to osteoblastic development (*Runx2*, *Bglap, Bglap2, Vdr, Sp7, Dmp1, Bmp3 and Pth1r*), but also included genes linked to epithelial to mesenchymal transition (*Notch3, Tbx2*), intercellular communication (*Cldn10, Cdh11*) and soluble factors of which CARc have not yet been described as a relevant source, such as angiotensinogen (*Agt),* Kininogens 1 and 2 *(Kng1, Kng2)* and Midkine *(Mdk)*.

In turn, the PαSc signature included genes characteristic of fibroblastic identity (Podoplanin [*Pdpn*], Endosialin [*Cd248*]) (Figure 2A and S2A), genes associated to angiogenic regulation (*Mfap5, Tnxb*) and cytokines (*Il33* and *Ccl11*). Given the poorly defined anatomical distribution of PαSc we assessed the specificity of some of the identified genes as putative markers of this subset. Flow cytometry analyses confirmed that within the CD45^−^Ter119^−^CD140b^+^ fraction, expression of Pdpn was strictly restricted to the PαSc, which homogeneously expressed high levels of this protein (98.9 ± 1.1 %) (Figure 2B). Using 3D imaging we observed a unique subset of Pdpn^+^Sca-1^+^CD140b^+^ pericytes, which exclusively localized in an adventitial layer of flattened stromal cells surrounding large, central BM arteries but not arterioles of smaller caliber (Figure 2C, 2D and Movie S1). Of note, bone-lining mature osteoblasts/osteocytes, which had not been included in our study, also expressed Pdpn but could be easily discriminated from PαSc by morphology, endosteal distribution and lack of expression of CD140b (Figure S2B and S2C). *Il33* expression was also exclusively detected in PαSc and strongly increased after stimulation with LPS (Figure S3A-D). In *Il33*-GFP reporter mice (Oboki et al., 2010), GFP^+^ cells were observed wrapped around large arteries, thereby confirming the adventitial localization of at least a fraction of cells categorized as PαSc (Figure S3D).

**Figure 2.**
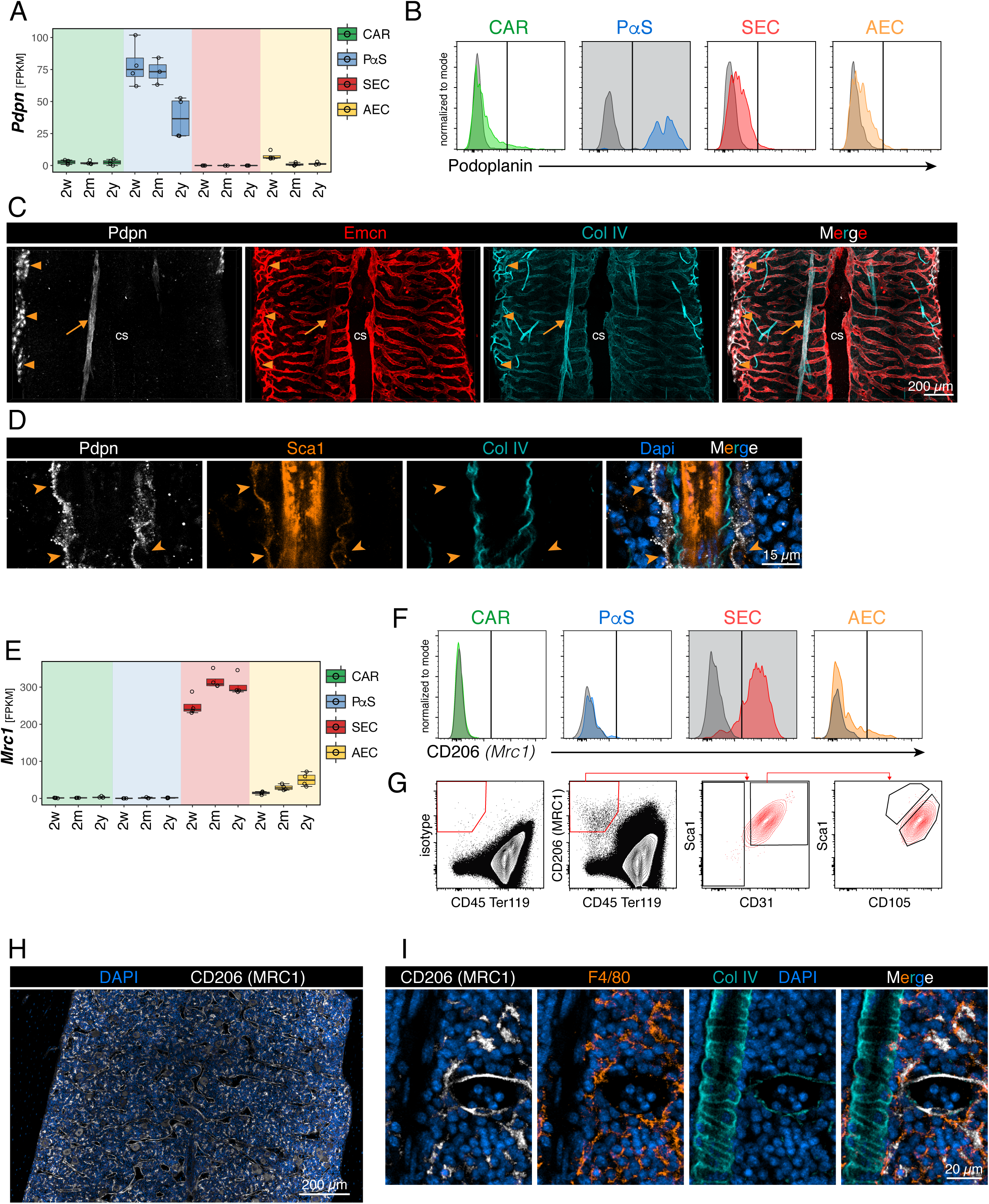
Podoplanin identifies periarterial PαS cells and sinusoids express distinctively high levels of mannose receptor, C type1. **A.** RNA-seq expression values (FPKM) for Pdpn (Podoplanin). **B.** Flow cytometry analysis of Pdpn expression by the four pre-gated stromal cell types. Isotype controls are displayed as grey histograms. **C.** Immunofluorescence staining of a thick femoral slice. Pdpn expressing cells are located close to the endosteal surface (bone-lining osteoblasts triangle shape) and around large arteries (PαSc arrow shape). Scale bar: 200 *µ*m. **D.** Higher magnification of a centrally located arterial region. The arrow heads point at peri-arterial Pdpn^+^ Sca1^+^ cells. Scale bar: 15 *µ*m. **E.** RNA-seq expression values (FPKM) for *Mrc1* in all cell types. **F.** Flow cytometry immunostaining of CD206 (*Mrc1*) for the four pre-gated stromal cell types. Isotype controls are displayed as grey histograms. **G.** Flow cytometry immunostaining of CD206 *(Mrc1)* displaying total BM. Pre-gated for singlets and live cells (DAPI^−^). The majority of CD206^+^, non-hematopoietic cells (CD45^−^ Ter119^−^) fall into the SEC gate. **H.** Immunofluorescence staining of CD206 in an adult BM diaphysis. Scale bar: 200 *µ*m. **I.** Higher magnification of a region showing representative examples of a large artery (CD206^−^), a sinusoid (CD206^+^) and macrophages expressing CD206, which are co-stained with F4/80 (orange). Scale bar: 20 *µ*m.

We next focused on selective gene signatures that allow straight forward separation of SECs from AECs. Among markers of AECs we found the established arterial specification gap junction genes *Gja4* and *Gja5* (encoding for connexin 37 and connexin 40), the transcription factor *Sox13* and the angiocrine factor *Jag2*. As for SECs, the gene signature included endothelial selectins (*Selp* and *Sele*), as well as three major scavenger receptors, namely Stabilin 1 and 2 (*Stab1, Stab2*) and the Mannose receptor-C type 1 (*Mrc1* also known as CD206, Figure 2E). The characteristic expression of phagocytic molecules in SECs underlie their potential role in mediating clearance of blood-borne particulate substances (X. M. Li et al., 2009). 3D imaging and FC confirmed that within the BM vascular network, CD206 expression was absent in arteries and arterioles, weakly emerged in TV and became very prominent in the entire sinusoidal tree (Figure 2F, 2G, 2H, 2I, S2D). CD206 was also expressed in a rather abundant population of BM cells identified as macrophages based on canonical marker expression (Figure 2I).

### Transcriptomic profiles reveal mesenchymal subset-specific functions in adult BM

To gain insight on the functional, ontogenic and molecular differences of the two mesenchymal populations under study, we performed global gene ontology (GO) analysis on the entire set of differentially expressed genes (DEGs) (log_2_ FC > 1, FDR ≤ 0.05). Consistent with the presence of both osteo- and adipo-primed progenitors in the CARc pool, genes upregulated in this subset were highly overrepresented in biological processes (BPs) related to bone morphogenesis, mineralization and endochondral ossification (Omatsu et al., 2010; Zhou et al., 2014)(Figure 3A, 3B), and included adipocytic factors such as *Adipoq* and *Pparg* (Figure 3B and 3C). In contrast, expression of osteoadipogenic markers was negligible or strongly downregulated in PαSc (Figure 3B and 3C), and GO categories overrepresented in the transcriptome of this cell type were related to the regulation of cartilage and connective tissue development (Figure 3A). Prototypical transcription factors and/or genes involved in chondrocyte specification such as *Sox9, Fmod, Prg4, Fxyd6 or Lum* were distinctively expressed at high levels by PαSc, which in contrast lacked expression of classical markers of mature chondrocytes (*Acan*) (Figure 3B) (Boeuf et al., 2008). Compared to CARc, PαSc showed distinctive expression of *Cd34*, which was confirmed at the protein level by flow cytometry (Figure 3C and 3D). Corroborating our previous imaging analysis, Pdpn^+^Sca-1^+^ adventitial cells ensheathing the main longitudinal femoral arteries expressed CD34 and low levels of CXCL12-GFP, thus corresponding to PαSc (Figure S4A-S4E).

**Figure 3.**
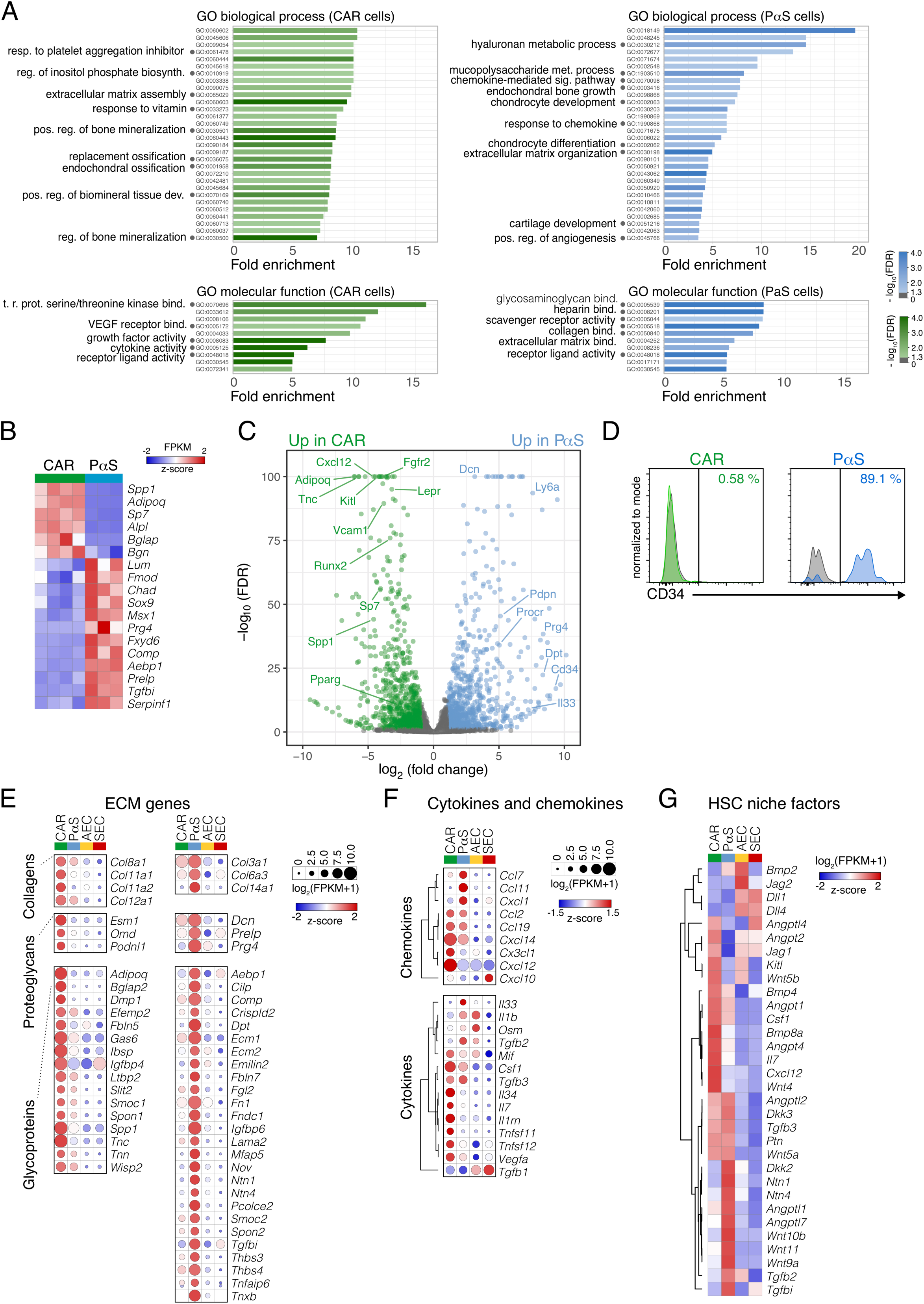
Transcriptomic comparison of BM mesenchymal cells highlights functional differences between CAR and PαS. **A.** Overrepresented gene ontology (GO) terms for genes differentially expressed in either CAR cells or PαS cells compared to the remaining three stromal cell populations (multigroup comparison, FDR ≤ 0.05, log_2_ FC ≥ 2). Full list of genes and DEGs is shown in Table S1. All displayed GO terms are significant (FDR ≤ 0.05). resp.: response, reg.: regulation, t.r.prot.: transmembrane receptor protein, pos.: positive. **B.** Heatmap of expression of classical osteoblastic or chondrocytic genes in CARc and PαS. Columns represent individual biological replicates. Row z-score normalized expression values. **C.** Direct pairwise comparison of DEGs between CARc and PαSc. Significant DEGs are highlighted in color (FDR ≤ 0.05, −1 ≥ log_2_ FC ≥ 1). Green dots: significantly up-regulated in CARc. Blue dots: significantly up-regulated in PαSc. **D.** Flow cytometry surface staining for CD34 on pre-gated CARc and PαSc. Grey histograms depict isotype control stainings. **E**. Heatmap of differentially expressed extracellular matrix (ECM) genes (FDR ≤ 0.05, log_2_ FC ≥ 2) from a multigroup comparison (CAR vs PαS & AEC & SEC and PαS vs CAR & AEC & SEC). All differentially expressed core-matrisome genes with high expression levels (FPKM ≥ 30) are displayed. Row z-score normalized median expression values. **F**. Row clustered heatmap of selected cytokines and chemokines with robust expression in at least one cell type. Row z-score normalized median expression values. **G.** Clustered heatmap showing expression of published HSC niche factors in all stromal cell types isolated from 2m old mice Row z-score normalized median expression values.

We found that the transcriptomes of both mesenchymal subpopulations were enriched in different aspects of extracellular matrix (ECM) biology, including production, assembly and binding to ECM components (Figure 3A). This was reflected in the abundant expression of a distinctive matrisomic signature of collagens, proteoglycans and glycoproteins in each mesenchymal subset (Figure 3E). ECM transcripts were also found in endothelial subpopulations, albeit to a much lower degree. These findings confirm the specific roles of mesenchymal cells in matrix production and remodeling, and strongly suggest a certain degree of cellular and topographical compartmentalization of the matrisome in the BM, which remains unexplored. CARc transcriptomes were enriched in gene categories related to cytokine and growth factor activity (Figure 3A), while PαSc, AECs and SECs subsets showed a much more restricted cytokine expression profile (Figure 3F). In CARc, we detected abundant transcripts of *Csf1* and *Il34*, the two known ligands of the macrophage colony stimulating receptor (M-CSFR), as well as basal expression of the prototypic T cell chemoattractant *Ccl19*, suggesting that CARc, or at least a fraction of this subset, may control basal trafficking of marrow-residing T cell subsets. Paradoxically, CARc not only expressed very high levels of *Cxcl12*, but also of *Cxcl14*, which has been proposed to function as a partial antagonist of CXCR4 function (Tanegashima et al., 2013). In line with their reported role in monocyte trafficking (Shi et al., 2011), CARc expressed *Ccl2* and *Cxc3cl1* in homeostasis. Finally, CARc were also most abundantly expressing a broad range of pro-hematopoietic growth factors (Figure 3G). Beyond *Cxcl12*, *Kitl, Il7, Angpt1 or Ptn* (Ding et al., 2012; Himburg et al., 2018; Sugiyama et al., 2006), we found substantial expression of *Bmp3* and *Wnt4* (Louis et al., 2008).

In turn, PαSc distinctively expressed certain chemokines and cytokines with known activities in myeloid cell migration and development (*Cxcl1*, *Ccl11*, *Ccl7* and *Il33),* the axon guidance protein *Netrin1 (Ntn1),* and a limited set of relevant pro-hematopoietic genes, which included the Wnt antagonists, *Dkk2* and *Dkk3*, as well as *Angptl1, Angptl2* and *Angptl7,* factors shown to expand human HSCs *in vitro* (Xiao, 2015; Xiao et al., 2015). Altogether, transcriptomic and spatial analyses suggest a potential, yet unrecognized niche regulatory activity of PαSc the HSC pool and hematopoiesis during adulthood.

### Major remodeling of stromal transcriptomic profiles in early postnatal stages

We next sought to elucidate sequential changes in gene expression patterns during postnatal life. As expected, we detected substantial transcriptional variations between 2w, 2m and 2y mice in all stromal subsets analyzed (Figure 4A). Strikingly, PCA and DEG analyses revealed broader modifications within the short period comprising the transition from juvenile to adult stages (45 days) than those arising in the progression from adult to aged stages (18 months difference) (Figure 4A). To obtain a refined and global view of temporal dynamic changes we employed Short Time-series Expression Miner (STEM) algorithms. Using this analytical tool, DEGs were grouped in age-dependent expression profiles that were then analyzed for enrichment in GO terms or pathways. In three out of four cell types (CARc, SECs and AECs) relevant clusters corresponded to DEGs that followed a continuous increase or decrease in expression during aging (designated cluster 1 (c1) and cluster 3 (c3), respectively) or DEGs that were significantly up- or downregulated from 2w to 2m old, and stabilized thereafter for the rest of the lifespan (grouped under c2 and c4, respectively Figure 4B). The most representative GO terms overrepresented in each cluster are shown in Figure 4B and a complete list is provided in Table S2.

**Figure 4.**
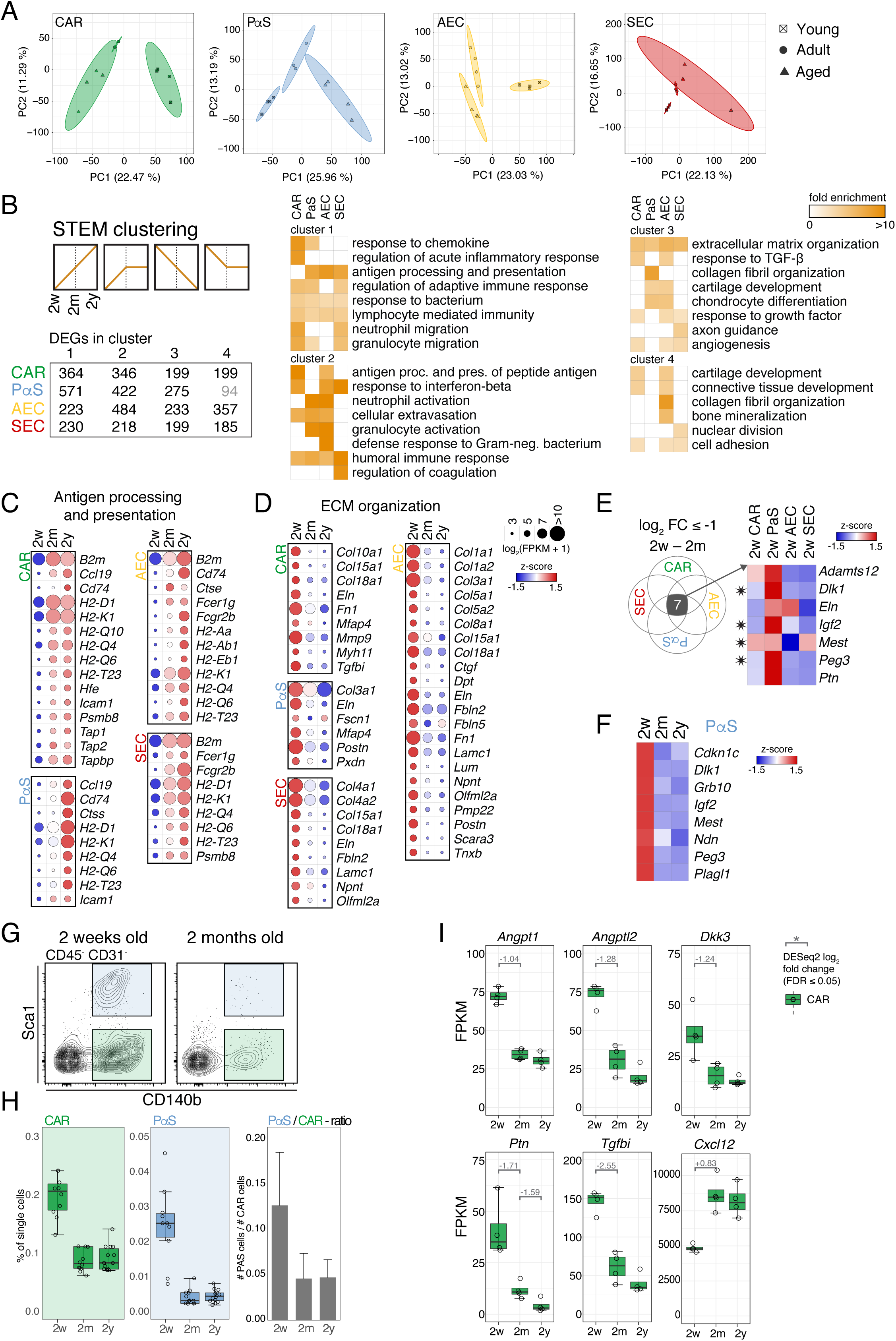
Global ontogenetic transcriptional changes. **A.** PCA of all three experimental conditions (2w, 2m, 2y) for each individual cell type (color coded). Analysis based on expression values of all detected genes. Individual dots represent biological replicates. **B.** Short time series clustering analysis using STEM (see experimental procedures). Clustering of DEGs from any inter-condition comparison into pre-defined expression profiles for each cell type. The four significantly overrepresented expression profiles in which DEGs cluster are schematically displayed. Bottom panel: number of DEGs allocated to each individual expression profile (black: FDR ≤ 0.05, grey: not significant, FDR ≥ 0.05). Heatmap displays fold enrichment of preselected overrepresented GO terms for each profile (full list of GO terms is provided in Table S2). Shown in orange color (independent of grading) are only significantly enriched categroies (FDR ≤ 0.05). **C.** Heatmap of genes belonging to the GO term antigen processing and presentation, which are significantly up-regulated (FDR ≤ 0.05, log_2_ FC ≥ 1) during the 2w to 2m transition. Row normalized expression values. Only genes with FPKM ≥ 30 are depicted. **D.** Heatmap of genes belonging to the GO term extracellular matrix (ECM) organization, which are significantly down-regulated (FDR ≤ 0.05, log_2_ FC ≤ −1) during the 2w to 2m transition. Only genes with FPKM ≥ 30 are depicted. **E.** Overlap analysis of DEGs (FDR ≤ 0.05, log_2_ FC ≥ 1 or log_2_ FC ≤ −1) downregulated during the 2w to 2m transition. Heatmap of the 7 down-regulated genes common to all cell types. Asterisk: genomically imprinted genes. Row normalized expression values. **F.** Heatmap of a selected imprinted gene network (Berg et al., 2011). Row normalized expression values. **G.** Flow cytometry scatterplot pre-gated for non-hematopoietic non-endothelial (CD45^−^CD31^−^) cells at 2 weeks and 2 months. Color coded CARc (green) and PαSc (blue) gate. **H.** Frequencies and calculated ratios of CARc and PαSc at all three experimental timepoints based on flow cytometry measurements. **I.** RNA-seq expression values for reported HSC niche factors that significantly change during the 2w to 2m timeframe. Values for log_2_ FCs from differential expression analysis are shown in grey inserted in the graph.

Since functional changes in the BM microenvironment during early postnatal development remain underexplored, we first focused in DEGs contained in c2 and c4, which were exclusively up- or downregulated in the juvenile to adult transition. Unexpectedly, c2 contained genes mostly associated to inflammation and immune activation in all cell types. For instance, genes related to antigen processing and presentation via major histocompatibility (MHC) class I, were highly upregulated in adult compared to juvenile CARc and SECs (Figure 4B and 4C). These included classical (*H2-D and H2-K1, H2-M3*), as well as less characterized non-classical MHC class I molecules (*H2-Q, H2-T*), genes related to peptide transport (*B2-m*, *Tap1, Tap2*) and antigenic presentation in the cell membrane (Figure 4C). Thus, similar to stromal lineages populating lymphoid organs, CARc and SECs are equipped with the molecular repertoire for antigen presentation, and the acquisition of this cellular machinery is developmentally timed to the entry into adulthood. Conversely, analysis of DEGs assigned to c4 revealed general overrepresentation of BPs related to ECM organization, cartilage and connective tissue development, cell adhesion, responses to transforming growth factor-β (TGF-β) and angiogenesis. Indeed, a number of genes encoding matrisomic proteins, ECM remodeling and degradation enzymes, glycoproteins, and adhesion molecules, were conspicuously downregulated in BM mesenchymal cells from 2w to 2m old BM (Figure 4D).

To further explore the nature of this developmental transition, we searched for DEGs that were concurrently downregulated in all stromal lineages in adult compared to juvenile BM. We found a unique signature containing only seven common genes, which includes *Ptn*, *Igf2* and *Dlk1,* genes that are known to play decisive roles in the regulation of self-renewing expansion of adult and embryonic(Chou and Lodish, 2010; Himburg et al., 2018; Kokkaliaris et al., 2016; Mascarenhas et al., 2009; Thomas et al., 2016) HSCs. Strikingly, all seven genes displayed by far the highest expression levels in juvenile PαSc compared to other stromal subsets, and the majority (four out of seven, *Igf2*, *Dlk1*, *Peg3* and *Mest)* belong to a group of genomically imprinted genes, which are monoallelicaly expressed in a parent of origin-specific fashion (Figure 4E). This prompted us to inspect the expression of a core subset of co-regulated imprinted genes, which are highly expressed in multiple somatic stem cells and downregulated in progenitors (Berg et al., 2011). We observed that this molecular fingerprint was abundantly expressed in juvenile PαSc, and sharply downregulated during progression to adulthood (Figure 4F). As found in previous studies, PαSc were significantly more abundant in juvenile than in adult BM tissues (Figure 4G and 4H). Altogether, these results suggest that PαSc display stem cell-like transcriptomic programs, are relatively more abundant in early postnatal BM, and express essential pro-expansive HSC cues, which gradually decline as adulthood is reached. Of note, in this temporal window, we also detected significant downregulation of HSC niche genes in CARc (*Angpt1*,*Angptl2*, *Dkk3* and *Ptn)*, whereas expression levels of *Cxcl12,* associated to HSC quiescence and maintenance followed the inverse trend (Figure 4I).

### Increased proinflammatory signatures in aged BM stroma

STEM analyses revealed that the enrichment in proinflammatory signatures was not exclusive of the transition from 2w to 2m, but was even more pronounced in 2y old mice (Figure 4B). A total of 274, 758, 273, and 159 genes were differentially expressed (log_2_ FC ≥ 1, FDR ≤ 0.05) in aged compared to adult counterparts CARc, PαSc, AECs and SECs, respectively. Using Gene set enrichment analysis (GSEA) we detected a general pronounced overrepresentation in upregulated genes of numerous inflammatory hallmarks and annotated REACTOME gene-sets, including cytokine and chemokine signaling, interferon (IFN) alpha and gamma responses and complement pathways (Figure 5A, 5B and 5C). Of major importance, aged CARc displayed increased expression of *Il1b* and *Il6,* key cytokines reported to enhance proliferation and myeloid bias, and impair HSC self-renewal during inflammatory processes (Pietras et al., 2016; Schürch et al., 2014). Among the most prominently upregulated genes in this subset were also a number of inflammatory chemokines (*Cxcl2, Cxcl5*), and most remarkably, several members of the complement cascade *(Cfd, Cfb, C4b, C3)* (Figure 5D). These data lend support to the notion that aging gradually results in an enhanced inflammatory milieu, at least partially driven by secretory activity of mesenchymal stromal cells, which drives alterations of BM hematopoiesis (Henry et al., 2015; Kovtonyuk et al., 2016). Strikingly, and in contrast to what has been proposed (Maryanovich et al., 2018), aged CARc did not exhibit changes in expression of crucial pro-hematopoietic factors such as *Cxcl12, Scf* or *Angpt1*. Nonetheless, we did note significant downregulation of Wnt-inhibitory factor (*Wif1*), *Angptl4* and *Npy,* associated to the modulation of HSC function (Figure 4I and 5E) (Park et al., 2015; Schumacher et al., 2015; Singh et al., 2017). Moreover, aging was linked to a repression of gene categories associated to ECM organization, collagen formation and adhesive interactions, which was strongest in mesenchymal cells but also evident in ECs. Multiple cell-specific matrisomic proteins that were highly expressed in CARc and/or PαSc from 2m old mice were downregulated in aged BM, including a number of collagens and enzymes involved in ECM formation (*Plod1, Plod2, Pcolce2*) and remodeling (*Mmp9, Mmp11*) (Figure 5F). Finally, functional annotation of DEGs in ECs during aging uncovered a significant upregulation of gene sets related to translation, posttranscriptional regulation and RNA and protein metabolism, concomitant with a decrease in angiogenic processes (Figure 5G and S5), which altogether are indicative of prototypic stress responses and could possibly be related to the disrupted arteriolar phenotype observed in aged BM (Maryanovich et al., 2018).

**Figure 5.**
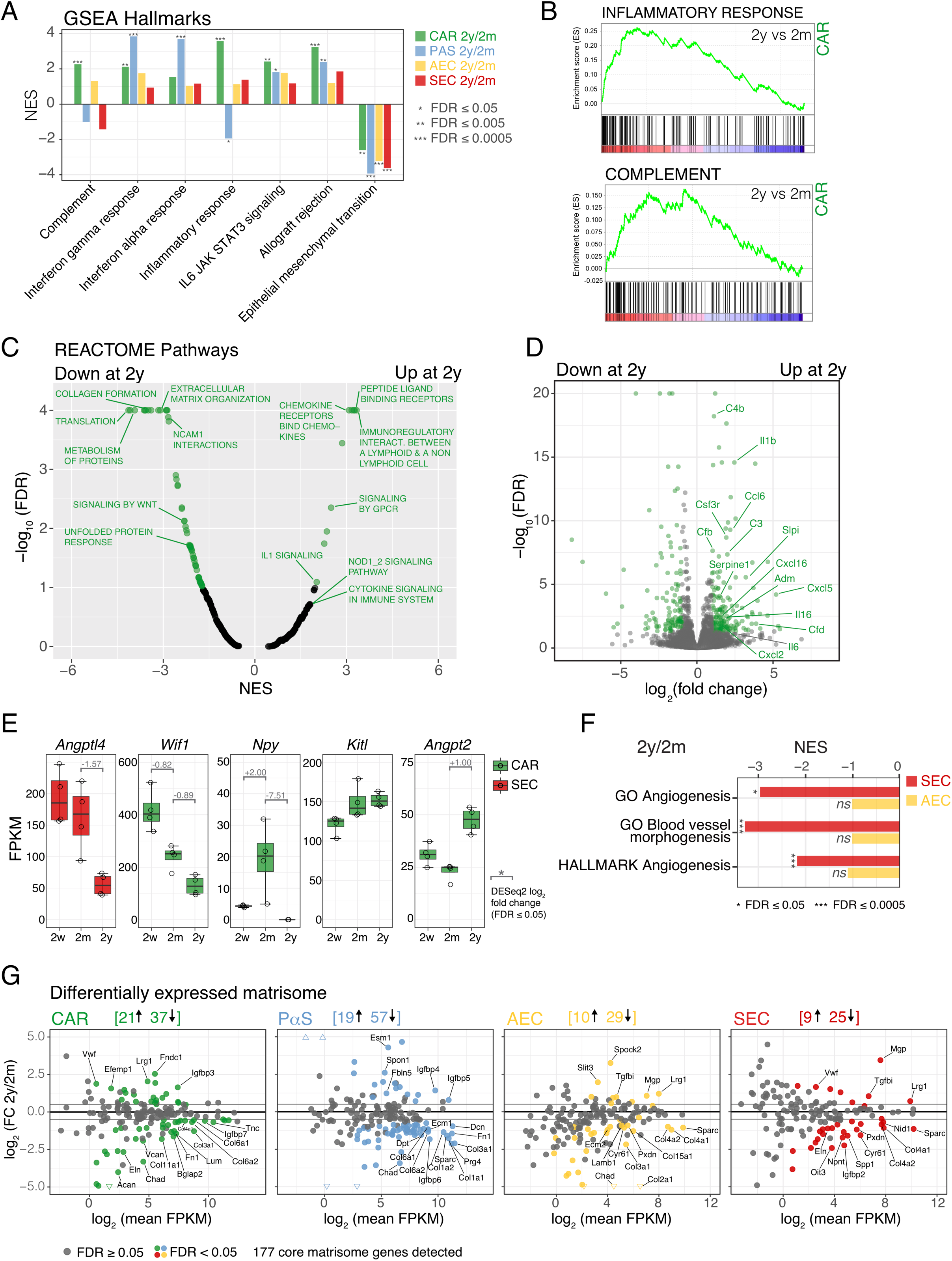
Basal inflammatory gene signature in aged BM stroma. **A.** GSEA results for selected significant hallmark gene sets. NES: Normalized enrichment score. **B.** GSEA enrichment plots for the gene sets inflammatory response and complement in CARc comparing cells isolated from 2y to 2m old mice (FDR ≤ 0.05). **C.** GSEA results for all REACTOME pathways comparing CARc from aged to adult mice. Significant gene sets are displayed in green color (FDR ≤ 0.05). **D.** Differential gene expression results for aged (2y) vs adult (2m) CARc. Significantly DEGs are highlighted in green (FDR ≤ 0.05, −1 ≥ log_2_ FC ≥ 1). **E.** Expression values for selected HSC niche factors. Grey numbers represent log_2_ FCs from differential expression analysis. **F.** GSEA results for angiogenesis related terms. **G.** Graphs depicting log_2_ FC of the pairwise comparison 2y vs 2m (Y axis) and log_2_ mean expression values (FPKM, X axis) for all core matrisome genes in each cell type. Significantly DEGs (FDR ≤ 0.05) are highlighted in color. Thick grey lines highlight log_2_ FC of −1 and 1 as a visual aid. Numbers on top of the plots indicate the total amount of DEGs that are up- or down-regulated.

### Overlap between aging and infection-induced transcriptional changes in BM stroma

Having established the proinflammatory nature of the aged BM microenvironment, we next compared the magnitude and quality of aging-induced expression changes, with those triggered acutely by administration of prototypical bacterial and viral infection-mimicking agents lipopolysaccharide (LPS) or polyinosinic-polycytydilic acid (pI:C). As previously described for HSPCs (Essers et al., 2009), the levels of Sca-1 were upregulated with variable intensities in all stromal populations during sterile inflammation (Figure S6A). Increases in Sca-1 expression led to the shift of a limited, yet significant fraction of SECs into the AEC gate, as well as of CARc into the PαSc gate, which partially contaminated sorted populations and rendered AEC and PαSc samples unsuitable for further analyses. Thus, we disregarded these subsets and exclusively focused subsequent data exploration on CARc and SECs. In general, short-term administration of both infection mimicking agents elicited profound transcriptional responses in both stromal cell types (Figure 6A, 6B and 6C, 6D). As expected, GSEA identified very strong overrepresentation of DEGs after LPS and pI:C in multiple pathways and hallmark sets associated to inflammatory processes and cytokine responses (Figure 6B). Enrichment in inflammation-related categories and overall changes in gene expression triggered by sterile infections were more pronounced than those observed in aged BM stroma and strongest in the case of pI:C (Figure 6C). Remarkably, we noted substantial overlaps in global expression trends and DEGs between LPS, pI:C and aging when compared to cells isolated from 2m old mice (Figure 6E). Indeed, out of 274 and 159 DEG upregulated in aged CARc and SECs (log_2_ FC > 1, FDR< 0.05), 117 (43%) and 56 (35%) were also found to be significantly increased in LPS and/or pI:C groups (Figure 6E). Among the commonly upregulated genes in two or more conditions we found chemokines, such as *Ccl5*, *Ccl6* and *Ccl19*, as well as *Cxcl9, Cxcl10* and *Cxcl11*, which are ligands of CXCR10 and whose expression is known to be strongly induced by IFN signaling. We also found increased expression of *Il6* in CARc of both LPS and pI:C treated mice, pointing once more to the potential preeminence of this factor in regulating functional aspects of inflammation-driven hematopoiesis. Interestingly, CARc massively increased expression levels of the IL1 receptor antagonist *Il1rn*, suggesting the activation of a protective mechanism to limit the deleterious effects of IL1 on hematopoietic cells (Figure 6F). In addition, gene sets related to apoptosis, hypoxia and RNA metabolism were also overrepresented in the transcriptome of stromal cells of mice treated with infection-mimicking agents (Figure 6B).

**Figure 6.**
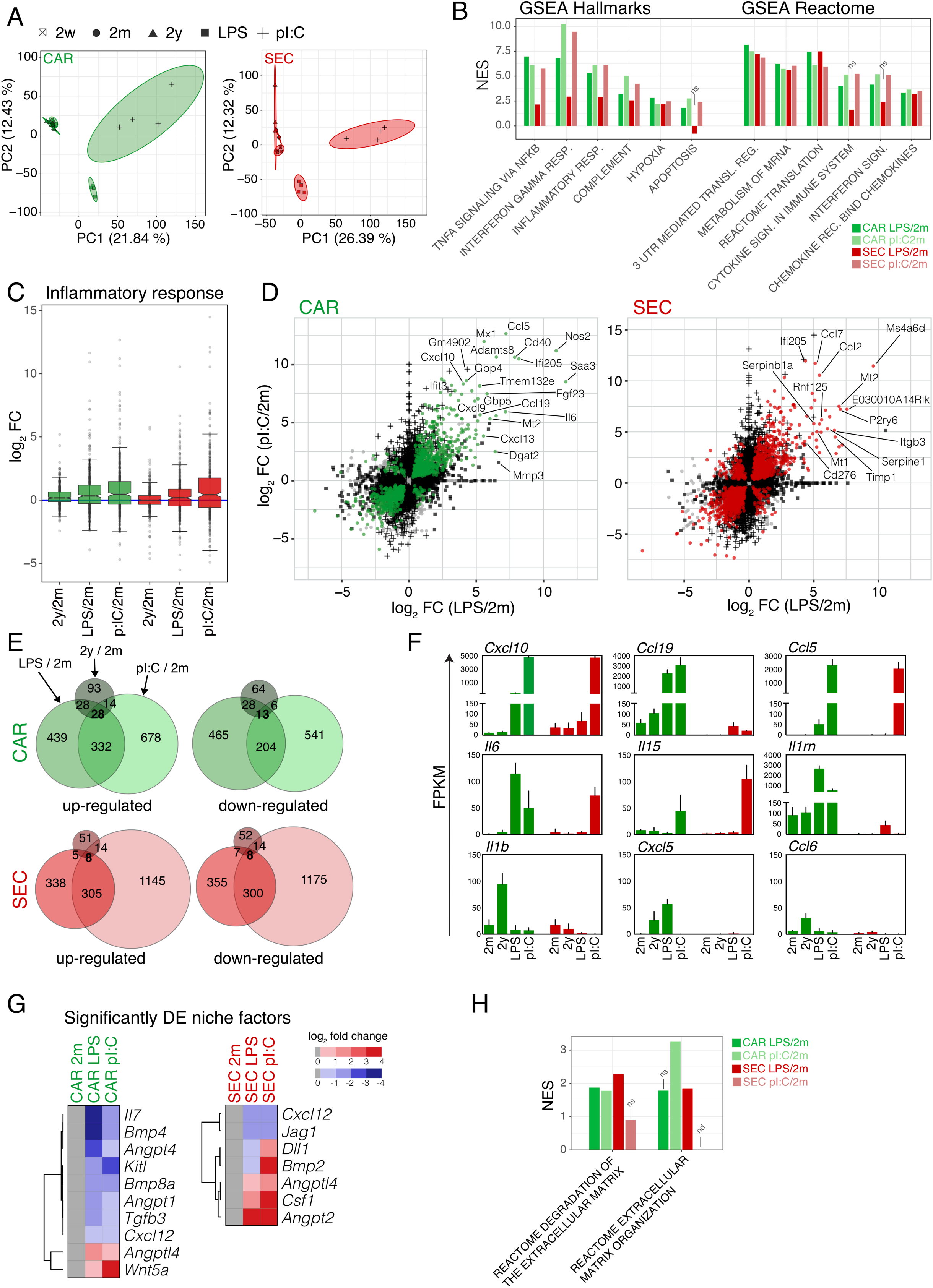
LPS and pI:C induce massive inflammation-associated transcriptional changes. **A.** PCA of all conditions including pI:C and LPS for CARc and SECs. **B.** Selection of significant gene set enrichment results for LPS and pI:C treatment in CAR cells and SECs. **C.** Log_2_ FC (indicated treatment vs adult control) of all genes belonging to the GO category inflammatory response for CARc (green) and SECs (red). **D.** Scatterplot displaying of log_2_ FC expression values of all genes in CARc and SECs in LPS (x-axis) and pI:C (y-axis) treatments compared to adult BM. Genes that are significantly differentially expressed (FDR ≤ 0.05) in one condition are shown as a black plus sign (pI:C/2m) or black square (LPS/2m), genes which are significant in both inflammatory conditions are displayed as colored dots (green or red) and not significant genes are shown in grey. Selected genes with large log_2_ FCs are highlighted. **E.** Overlap analysis of DEGs (FDR ≤ 0.05, log_2_ FC ≥ 1 or log_2_ FC ≤ −1) for the conditions: LPS, pI:C and 2y with respect to adult BM. **F.** Expression patterns of selected cytokines and chemokines. **G.** Heatmap of differentially expressed hematopoietic niche factors displaying color-coded log_2_ FCs (LPS vs 2m and pI:C vs 2m) for CARc and SECs. All shown genes are significant (FDR ≤ 0.05). **H.** GSEA results for significant REACTOME gene sets related to extracellular matrix (FDR ≤ 0.05). ns: not significant (FDR ≥ 0.05). nd.: not determined.

Other conspicuous changes were related to the hematopoietic supportive activity of stromal cells. Administration of LPS and pI:C led to an evident decline in the hematopoietic supportive function of CARc, through the strong downregulation of the most characteristic genes involved in extrinsic control of lymphoid and myeloid and HSPC maintenance specification (*Cxcl12, Kitl, Il7)* (Figure 6G). In contrast, we detected increased expression of *Wnt5a*, a niche-derived factor required for engraftment of functional HSCs upon transplantation in myeloablated recipients (Schreck et al., 2017). Similar to CARc, expression of pro-hematopoietic genes in SECs was also highly reactive to infection mimicking agents (Figure 6G). Finally, inflammatory stimuli strongly targeted matrisomic gene expression. Major ECM genes, but specially enzymes related to degradation and remodeling of ECM matrices in CARc and SECs were differentially expressed in LPS and pI:C treated BM with respect to adult controls (Figure 6H & S6B). These effects are potentially linked to a rapid tissue remodeling activity needed to reroute BM hematopoietic output under infections. In summary, our results highlight the dynamic responsive nature of BM stromal cells to infections and/or ensuing inflammatory processes, and the potential induction of stress gene expression programs in the context of an inflamed and aged BM microenvironment.

## Discussion

Crucial aspects of age-related changes in BM hematopoiesis and HSC function have been elucidated through comprehensive transcriptional, epigenetic and metabolic studies (Chambers et al., 2007a; 2007b; Gazit et al., 2013; Sun et al., 2014). In contrast, how BM stromal cell function evolves during physiological aging, and whether stromal perturbations correlate with changes in hematopoietic function remains underexplored. In this study we aimed to dissect novel features of previously described stromal populations and investigate fundamental changes in the molecular make-up of the BM microenvironment throughout postnatal life. We performed bulk RNA-seq of highly pure cell populations that were sorted using validated phenotypic signatures. This approach has the obvious disadvantage of neglecting the heterogeneity that is known to exist within many cell populations, which can only be captured using single cell (sc) transcriptomic technologies, widely employed nowadays. Nonetheless, based on recently published sc-RNAseq data, out of the four populations studied here, CARc are the sole fraction that has been shown to comprise a highly diverse set of subpopulations, while AECs, SECs and possibly PαSc, constitute largely homogeneous subsets (Baryawno et al., 2019; Tikhonova et al., 2019). Therefore, we here opted for population-based transcriptomic analyses, which afford deeper molecular insight and allowed us to resolve and compare stromal transcriptional landscapes with highest resolution. Perhaps most importantly, our experimental approach required the isolation of pure cellular subsets following protocols, which involve mechanical disaggregation and cell extraction processes, even though our most recent work demonstrates that substantial fractions of CARc, SECs and most likely other stromal populations are lost during such procedures. This represents an obvious potential shortcoming that nevertheless, equally applies to all studies on BM stroma reported to date, independently on the sequencing technology used, and therefore needs to be acknowledged when interpreting results. Having noted this, the valuable datasets here presented i) provide molecular fingerprints that inform on potentially novel stromal subset-specific functions and niche affiliations, ii) reveal the previously unappreciated dynamic molecular remodeling of the BM microenvironment during the postnatal window, and putative factors supporting the self-renewing expansion of HSCs detected in this phase iii) uncover the prominent activation of inflammatory transcriptional programs in stromal cells during aging, which resemble those elicited by infections and are mainly characterized by abnormal expression of pro-hematopoietic factors, cytokines and extracellular matrix-related genes.

We exploit cell-specific molecular fingerprints to infer relevant unknown functional aspects and unequivocally trace the anatomical localization of PαSc. Originally defined as a population enriched in mesenchymal stem cell potential, this cell type has been mostly investigated through the use of *in vitro* cultures, orthotopic or systemic transplantation (Hu et al., 2016; Nusspaumer et al., 2017), and their principal biological features remain undefined. We unexpectedly observed that PαSc express a signature associated to pre-chondrogenic potential, which stands in contrast to the strong osteoblastic and adipogenic signatures of CARc. Although previous work suggested that PαSc could reside in the proximity of arteries and arterioles (Morikawa et al., 2009; Omatsu et al., 2014), we have precisely mapped PαSc to the external adventitial layer of the large, centrally running femoral arterial branches, which is absent in smaller arterioles. In periarterial bundles, PαSc lie adjacent to, but are distinct from smooth muscle actin cells and *Nestin*-GFP cells (not shown) and can be distinguished by expression of CD34. Altogether these findings lead us to speculate that PαSc potentially represent a distinct lineage of mesenchymal stromal precursors equivalent to those recently found in the adventitial cuffs of multiple organs (Sitnik et al., 2016). Of interest, recent work suggests a central role of these CD140b^+^Sca-1^+^Pdpn^+^ in the maintenance of type 2 innate lymphoid cell (ILC-2) within defined periarterial niches of lung and adipose tissues via expression of IL-33 (Dahlgren et al., 2019; Mahlakõiv et al., 2019; Spallanzani et al., 2019). Given the strong anatomical and molecular analogies between these cells and BM PαSc, their potential contribution to ILC-2 developmental niches in the BM deserves to be explored. Beyond this, PαSc have been suggested to contribute a critical fraction of CXCL12 production for HSC maintenance (Greenbaum et al., 2013). However, whether and to what extent this subset influences early stages of hematopoiesis is unclear. We find that in adult mice, PαSc express a distinct set molecular factors which mainly include key HSC pro-expansive factors such as *Angptl2, Angptl4*. Thus, given their restricted spatial localization and cytokine expression profile, we anticipate that this cell type fulfills unknown functions in hematopoietic control, which need to be addressed through the development of tailored models allowing targeting of PαSc cells and their lineage tracing *in vivo*. The distinctive PαSc gene signature here identified could prove valuable for the rational design of such experimental tools.

Unlike PαSc, a large number of studies have exposed multiple functional facets of CARc. Our data suggest that CARc as a whole achieve this remarkable functional pleiotropism through expression of a large plethora of different chemokines and cytokines that potentially regulate not only ontogeny, but also trafficking of myeloid and lymphoid populations and immune function of the BM during homeostasis. For instance, CARc abundantly express *Il34* and *Csf1*, suggesting yet another previously unappreciated function of this cell type in controlling macrophage development and maintenance. These data reinforce the existence of an intimate bidirectional functional relationship between CARc and macrophage networks (Casanova-Acebes et al., 2013; Chow et al., 2011). Future investigations will shed light on whether this, as well as the multiple other jobs of CARc in hematopoietic regulation are independently performed by the recently described CARc subtypes in distinct anatomical niches (Baryawno et al., 2019; Tikhonova et al., 2019).

While much attention has been casted on the study of aging and BM function, the mechanisms underlying the defining features of juvenile hematopoiesis and its progression to adult stages remain underexplored. Neonatal BM tissues are comparatively much more permissive to engraftment of embryonic HSCs than adult counterparts, which points to the supply of unique extrinsic regulatory signals by the BM microenvironment at these stages (Arora et al., 2014). In line with this, our data demonstrate that, despite conserving phenotypic traits, the four stromal cell types investigated display substantial modifications in their transcriptional landscape with respect to adult stroma. These differences, which are most likely linked to the pronounced tissue remodeling activity during organ growth, may also account for the active cell-extrinsic orchestration of continuous HSC proliferation in the postnatal window. Remarkably, although embryonic emergence and production of HSCs have been observed to depend on low-grade proinflammatory cues in hematopoietic tissues (Espin-Palazon et al., 2018; He et al., 2015; Sawamiphak et al., 2014; Trompouki, 2016), we find that cells of the stromal microenvironment in juvenile mice are almost completely devoid of expression of all inflammatory mediators, MHC class I molecules and IFN responsive genes, whose expression is relatively abruptly turned on as adulthood is reached. Although we cannot rule out that at this point basal proinflammatory signals are provided by BM-resident innate immune cells not analyzed here, our data strongly suggest that postnatal HSC expansion most likely takes place in a virtually sterile BM milieu (Adkins et al., 2004). We hypothesize that the non-inflammatory nature of juvenile BM tissues could represent a safeguard mechanism to protect HSCs from the deleterious consequences of uninterrupted proliferation and increased cumulative divisional history, as observed in inflammation-induced HSC cycling (Takizawa et al., 2017; Walter et al., 2015). In fact, little is known to date on the specific niche-derived signals that promote self-renewing expansion during juvenile development. We found that PαSc in these stages express a distinct set of cytokines and adhesion molecules with reported properties in HSC expansion, which are rapidly down-regulated during the transition to adulthood. Among them, IGF2 is a mitogenic factor for HSCs, which is produced by putative niche cell types of the fetal liver (Chou and Lodish, 2010; Mascarenhas et al., 2009; Thomas et al., 2016; Wu et al., 2008). Deregulation of the Igf2-IGFR1 signaling axis in adult BM leads to impaired quiescence, uncontrolled proliferation and eventual exhaustion of the HSC compartment in adults (Venkatraman et al., 2013). Juvenile PαSc also expressed very high levels of *Dlk1*, *Dpt* and *Ptn*, all of which have been assigned HSC supportive activity *in vitro* and/or *in vivo* (Kokkaliaris et al., 2016; Wu et al., 2008). Given these marked changes in expression profiles and that PαSc are relatively much more abundant in juvenile than in adult BM, it is likely that this cell-type provides core, developmental-stage specific cues for temporary HSC expansion. The identification of the principal common molecular signals promoting HSC self-renewing proliferation in fetal liver and juvenile BM could hold the key for the development of efficient protocols for HSC expansion *ex vivo*.

Our work further sheds light on the fundamental impact of aging on stromal cell biology and BM hematopoiesis. The emergence of a strong proinflammatory signature in the stromal compartment, and particularly in mesenchymal cells, is in line with previous findings on thymic stroma (Ki et al., 2014), and demonstrates that deleterious age-related effects are not limited to the parenchyma of organs, but similarly inflicted on stromal infrastructures, whose contribution to aging phenotypes in tissues is very relevant. Unexpectedly, we did not detect extensive or substantial changes in expression of principal factors regulating hematopoiesis, in CARc. Thus, most likely, aging alterations in the stromal control of hematopoiesis are mediated through upregulation of multiple cytokines and chemokines. Among many others, we identify *Il1b* to be massively increased in CARc from aged mice. Given its recently reported activities in trained immunity, promotion of myelopoiesis and restriction of self-renewal of HSCs, stromal derived IL-1 is a prime niche candidate for driving age-related HSC deficits, clonal hematopoiesis and its vascular manifestations (Chavakis et al., 2019; Pietras et al., 2016). The induction of a basal inflammatory context is also reflected in the upregulation of multiple components of the complement system, which so far had been associated to HSPC mobilization to peripheral tissues during stress hematopoiesis (Ratajczak et al., 2010). Overexpression of complement genes is a hallmark of aging tissues and abnormal activation of this system has been linked to the emergence of degenerative diseases and cancer (de Magalhães et al., 2009; Komuro et al., 2012; McGeer et al., 2005). Thus, it will be crucial to understand to what extent complement cascades and the cytokines here identified induce a selective pressure that promotes aging-induced clonality and contribute to increased incidence of hematologic neoplasia (Henry et al., 2011; Vas et al., 2012). Our analyses reveal that the transcriptomic signatures activated during aging, partially resemble those induced by both infection-mimicking agents pI:C and LPS employed here. In aged CARc and SECs we found overexpression of prototypical genes downstream of IFN, as well as TLR4 signaling axes. Of note for frequently used experimental approaches, the vast and rapid changes in the transcriptome observed upon pI:C stimulation should bring caution to studies employing the *Mx1-*Cre system in which repeated administration of this agent is used to delete genes in both hematopoietic and stromal compartments. Another remarkable hallmark of aged BM stroma, similarly recapitulated during sterile infections, is the generalized reduction of ECM related genes and, most strongly, of different collagens, which has been also observed in fibroblasts of other organs (Salzer et al., 2018). Matrisomic properties of the surrounding microenvironment are known to exert major influences in cell fate decisions in BM mimicking 3D cultures (Choi and Harley, 2017). However, teasing apart the specific biomechanical and molecular effects of these structures *in vivo* has been so far challenging due to the daunting complexity of matrix fiber scaffolds. Our data suggest that ECM composition is i) largely derived from mesenchyme but molecularly compartmentalized between cellular subsets ii) very dynamically and specifically shaped not only in aging, but also throughout postnatal development and as part of rapid stereotypic responses to inflammatory challenge. It is therefore tempting to speculate that matrisomic scaffolds represent a malleable and actionable platform for the swift modulation of hematopoiesis by stromal cells in different physiological and pathological contexts. Future studies on ECM will require the combination of novel genetic tools and high-resolution quantitative imaging technologies for structural and biophysical analyses.

In summary, the temporally-resolved, population-based, transcriptomic profiles of stromal subsets presented here, represent powerful resources for the dissection of unappreciated functional aspects of stromal cells, and the understanding of changes in stromal-hematopoietic crosstalk that lead to modulation of BM function throughout the lifespan of an organism.

## Methods

### Mice

C57BL/6JRj mice were purchased from Janvier Labs (France) and maintained under standard conditions at the animal facility of the University Hospital Zurich. *Il33*-GFP mice were generated by the RIKEN Center for Life Science Technologies (Kobe, Japan) (Oboki et al., 2010). All the experimental procedures involving animal models were approved by the veterinarian office of the Canton of Zurich, Switzerland.

### Experimental conditions

Male animals were used for RNA-seq experiments. Mice were analyzed at different ages, namely 2 weeks (14-15 days old, termed juvenile), 2 months (8-9 weeks old, designated as adult), 2 year-old (20-24 months old, aged mice). LPS-challenged mice (8-9 weeks old) were injected twice intraperitoneally with 35 µg of lipopolysaccharides (in 200 µL phosphate buffered saline). pI:C-challenged mice (8-9 weeks old) were injected twice intraperitoneally with 100 µg of polyinosinic-polycytidylic acid (in 200 µL phosphate buffered saline). The injections were performed 48 hours and 4 hours before sacrifice. Four independent biological replicates, which were collected at different times and dates were processed and sequenced for each experimental group. All replicates, cell types and conditions were sequenced at once.

### Stromal cell isolation

The isolation procedure for stromal cells has been previously described in detail in (Gomariz et al., 2018). Briefly, two sets of femur/tibia/pelvis were isolated from the surrounding tissue and cleaned using a surgical scalpel and paper tissues in order to thoroughly remove surrounding muscle and connective tissue. The BM content was flushed out into 6-well plates utilizing a 26G syringe and 5 mL of digestion medium (DMEM GlutaMAX™, 10 mM HEPES, 10 % fetal bovine serum). The remaining bone enclosures were carefully cut into small fragments (approximately 1 mm^3^ in size) using scissors and added to the same well already containing digestion medium and the previously flushed marrow. For the enzymatic tissue digestion, collagenase (0.04 g/mL) and DNAse (0.2 mg/mL) were added and thoroughly mixed using a 1 mL pipette and the cell suspension incubated for 45 minutes at 37 °C with gentle sample agitation. Next, 5 mL of ice-cold calcium- and magnesium-free phosphate-buffered saline (PBS) containing 10 % FBS were added and the suspension filtered through a 70 *µ*m cell strainer and washed once more with PBS (4 °C).

### Cell sorting and flow cytometry analysis

Cell suspensions were blocked using TruStain fcX^TM^ for 15 mins at 4 °C and subsequently immunostained using pre-conjugated surface antibodies for 30 minutes at 4 °C. All antibodies employed in this study, including commercial sources and concentrations are shown in Table S3. After staining, the cells were washed twice using ice-cold PBS and resuspended in PBS containing DAPI (0.5 μg/mL) and analyzed using an LSR II Fortessa (BD Biosciences). Post-acquisition data analysis was done using FlowJo 10 software. Cell sorting was performed using a FACS Aria (BD Biosciences) and all four stromal cell populations were sorted directly into RNase-free microfuge tubes (ThermoFisher) containing RLT lysis buffer (Qiagen) at 4 °C until 4’000 to 50’000 cells were collected for each cell type. In order to collect enough cells, the cell extracts of two mice were combined for each replicate before cell sorting.

### RNA isolation, library preparation, cluster generation and sequencing

Following the manufacturer’s instructions of the RNeasy Plus Micro Kit (Qiagen), genomic DNA was depleted using gDNA eliminator columns and RNA extracted from the cell lysates. The quality of the isolated RNA was measured using a Bioanalyzer 2100 (Agilent, Waldbronn, Germany). Library preparation, cluster generation, and sequencing were performed at the functional genomics center Zurich (FGCZ, Zurich, Switzerland). The libraries were prepared using the SMARTer® Stranded Total RNA-Seq - Pico Input Mammalian - kit (Takara Bio, USA) following the manufacturer’s instructions. For each sample 600 pg of input total RNA were used. The TruSeq SR Cluster Kit v4-cBot-HS (Illumina, Inc, California, USA) was employed for cluster generation using 8 pM of pooled normalized libraries on the cBOT. Sequencing was performed on the Illumina HiSeq 4000 single end 125 bp using the TruSeq SBS Kit v4-HS (Illumina, Inc, California, USA).

### Bioinformatic analyses

The RNA sequencing dataset has been deposited and can be accessed under GSE133922. Processing of RNAseq data was done using the Nextpresso pipeline v.1.9.2 (Graña et al., 2018). The quality of the raw data was assessed using FastQC v0.11.3 (Andrews, 2010) and FastQ Screen v0.5.2 (Wingett and Andrews, 2018) tools. Cutadapt (Martin, 2011) was used to remove the adaptors, trim the first three bases of the reads and the 3’ and 5’ end nucleotides with a quality cutoff of 20, requiring a minimum length of 40. Reads were then aligned with TopHat 2.0.10 (Trapnell et al., 2012)using Bowtie 1.0.0(Langmead et al., 2009) and SAMtools 0.1.19 (H. Li, 2011) against the GRCm38/mm10 assembly of the mouse genome containing only protein coding genes. During the alignment only 2 mismatches and 20 multihits were allowed. Transcripts were quantified using htseq-count from HTSeq framework 0.6.1 (Anders et al., 2015) and the differential expression test was done with DESeq2 v1.18.1 (Love et al., 2014) using the Mus musculus GRCm38/mm10 transcript annotations from https://ccb.jhu.edu/software/tophat/igenomes.shtml. Fragments Per Kilobase of exon per Million fragments mapped (FPKM) expression values were computed using the Cuffquant and Cuffnorm functions included in Cufflinks V2.2.1 suite (Trapnell et al., 2012). Only genes showing enough global expression and variability were kept (> 2 FPKM in > 20% of the samples, IQR range > 1).

To check for similarities between replicates a Pearson correlation test was applied over the normalized and regularized logarithmic counts obtained from DESeq2. Only one sample displayed a correlation coefficient smaller than 0.90 in a comparison to its replicates and was therefore excluded from the analysis.

Principal Component Analysis (PCA) and Hierarchical Clustering were created using R (v3.4.1) and the ggpubr and dendextend packages over the normalized FPKM expression values. The Hierarchical Clustering used the Eucledian distance and the Ward.D2 algorithm as the clustering method.

Gene set enrichments were calculated using the GSEA software (Broad Institute) using the previously defined genes sets from the Molecular Signatures Database (MSigDB 6.2) (Mootha et al., 2003; Subramanian et al., 2005). The gene list was ranked according to log_2_ FCs from the differential expression test and the pre-ranked GSEA was run using the classic enrichment statistic with 10’000 permutations.

Gene ontology analysis was performed using the PANTHER Overrepresentation Test (GO database release: 2018-04-04). For each test a background list containing all detected genes in this study was provided and statistics evaluated using Fisher’s Exact test with FDR multiple test correction (Mi et al., 2018). Additionally, larger GO categories were selected by excluding GO terms that contained less than 10 genes in the background list.

The Short Time-series Expression Miner (STEM) has been previously described (Ernst et al., n.d.). Briefly, the input data table contained median FPKM expression values of all DEGs from any inter-condition comparison (2w/2m, 2w/2y, 2m/2y, FDR ≤ 0.05, −1 ≥ log_2_ FC ≥ 1). The following parameters were utilized: log-normalization of the data, STEM clustering method, a maximum unit change in model profiles of 2, and a permutation test including timepoint 0 calculating all possible permutations. The integrated GO analysis tool was used. A background list of all detected genes in this sequencing data set was provided, a minimum GO level of three selected, the minimum number of genes within a GO category set to five, and multiple hypothesis correction by randomization used. Heatmaps are based on the FPKM expression values if not stated otherwise and created using the online tool Morpheus (https://software.broadinstitute.org/morpheus).

### Immunostaining and volumetric confocal imaging

3D imaging protocols have been previously described in detail (Gomariz et al., 2018). Briefly, femurs were isolated and thoroughly cleaned from the surrounding tissue, fixed in 2 % paraformaldehyde in PBS (6h, 4°C), dehydrated in 30 % sucrose in PBS (72h, 4°C), embedded in OCT medium, snap-frozen and bi-laterally sectioned using a cryotome. The obtained thick BM slices were incubated in blocking solution overnight (0.2 % Triton X-100, 1 % bovine serum albumin, 10 % donkey serum, in PBS) at 4°C and subsequently stained with primary and secondary antibodies diluted in blocking solution (Table S3) for 72 hours each, including a 3 x 1 hour washing step (PBS) in between. The samples were optically cleared in RapiClear 1.52 by immersion for 6 hours. Imaging was performed on a SP8 Leica confocal microscope. Imaris software (v8.41) was used for analysis and rendering of microscopy data.

The authors declare no conflict of interest

## Author contributions

P.M.H. designed and performed experiments, analysed data and wrote the manuscript. R.G designed and performed experiments. S. B participated in the design of the study. EP-Y and F.A designed the bioinformatic pipelines and performed data analysis. M.G.M and C.N-A. contributed equally, designed and directed the study and wrote the manuscript.

## Acknowledgements

We thank Daniel Pinschewer (University of Zurich) for providing the *Il33-Gfp* knock-in mice. P.M.H is supported by fellowship from the Forschungskredit of the University of Zurich. This work was supported by a grant from the Swiss National Science Foundation to M.G.M (310030B_166673/1) and grants from the Swiss National Science Foundation(31003A_159597/1), FP7 Marie Curie Career Integration Grant (PCIG13-GA-2013-618633) from the European Union, and the Helmut Horten Foundation (Lugano, Switzerland) to C.N.-A.

---

**Figure S1. Related to Figure 1.**
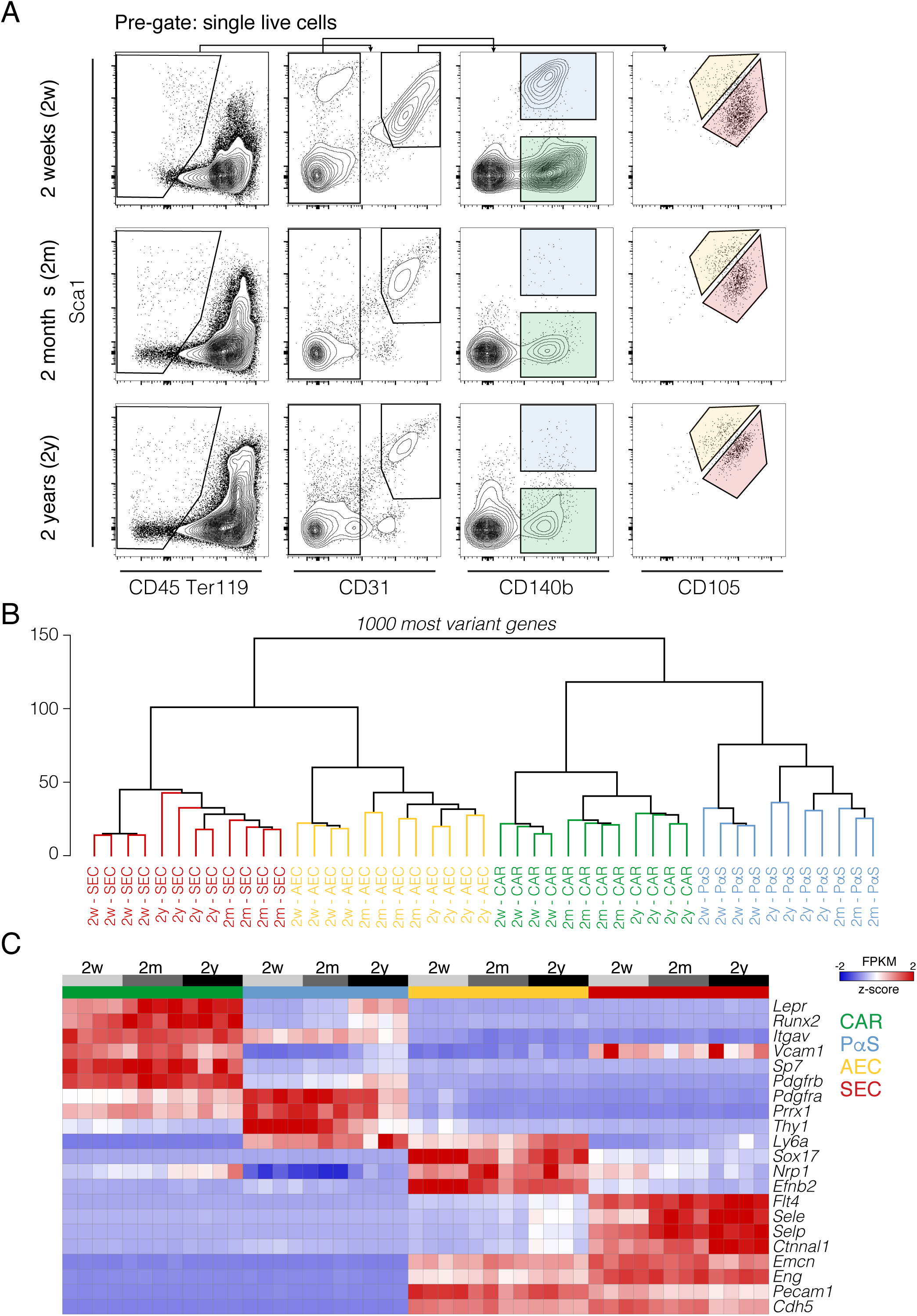
**A**. FACS gating strategy of BM stromal cell populations for all three experimental time-points. Pre-gated for singlets and live cells (DAPI-). **B**. Unsupervised hierarchical clustering of all cell types, timepoints and replicates using the top 1000 most variant genes. **C.** Heatmap of known stromal cell lineage specific markers for all timepoints. Row normalized expression values. Replicates are included as separate columns.

**Figure S2. Related to Figure 2.**
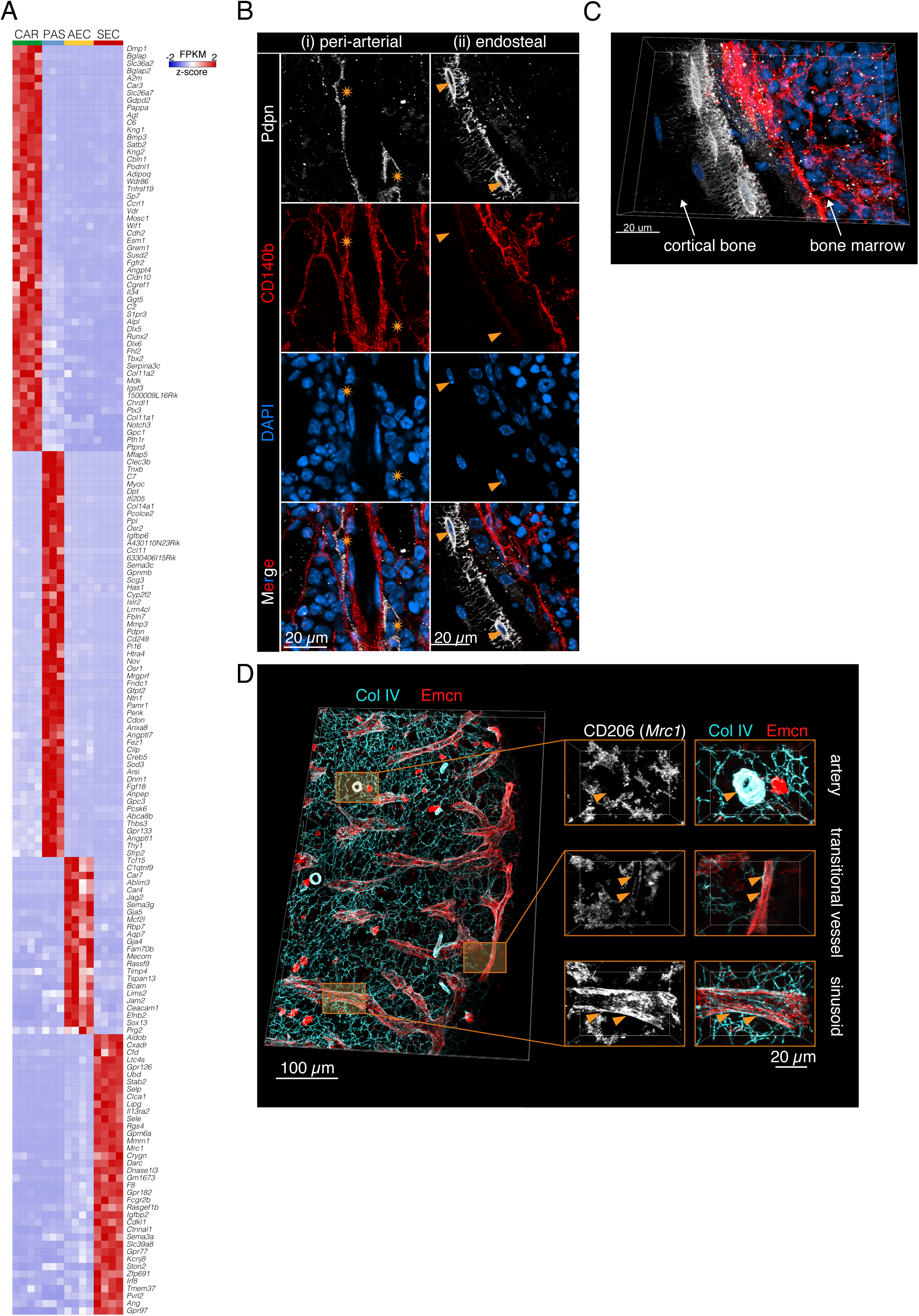
**A**. Heatmap of genes constituting highly cell type-specific markers from a multigroup differential expression analysis for the adult condition. Genes were filtered using the following criteria: FC “cell type of interest” vs “all other cell types” ≥ 10 (log_2_ FC ≥ 3.322), median FPKM (“cell type of interest”) ≥ 30, median FPKM (“all other cell types”) ≤ 30. **B.** Immunofluorescence co-staining of Pdpn and CD140b. Asterisks: Peri-arteial Pdpn^+^ cells. Arrow heads: Endosteally located Pdpn^+^ cells. Scale bars: 20 *µ*m. **C.** Volumetric representation of B (ii), high-lighting the morphology of endosteally located Pdpn+ cells. Scale bar: 20 *µ*m. **D.** Volumetric representation of a transversally cut BM diaphyseal slice immunostained for endothelial (Emcn), ECM (Col IV) and CD206. Scale bars: 100 *µ*m (left), 20 *µ*m (right).

**Figure S3. Related to Figure 2.**
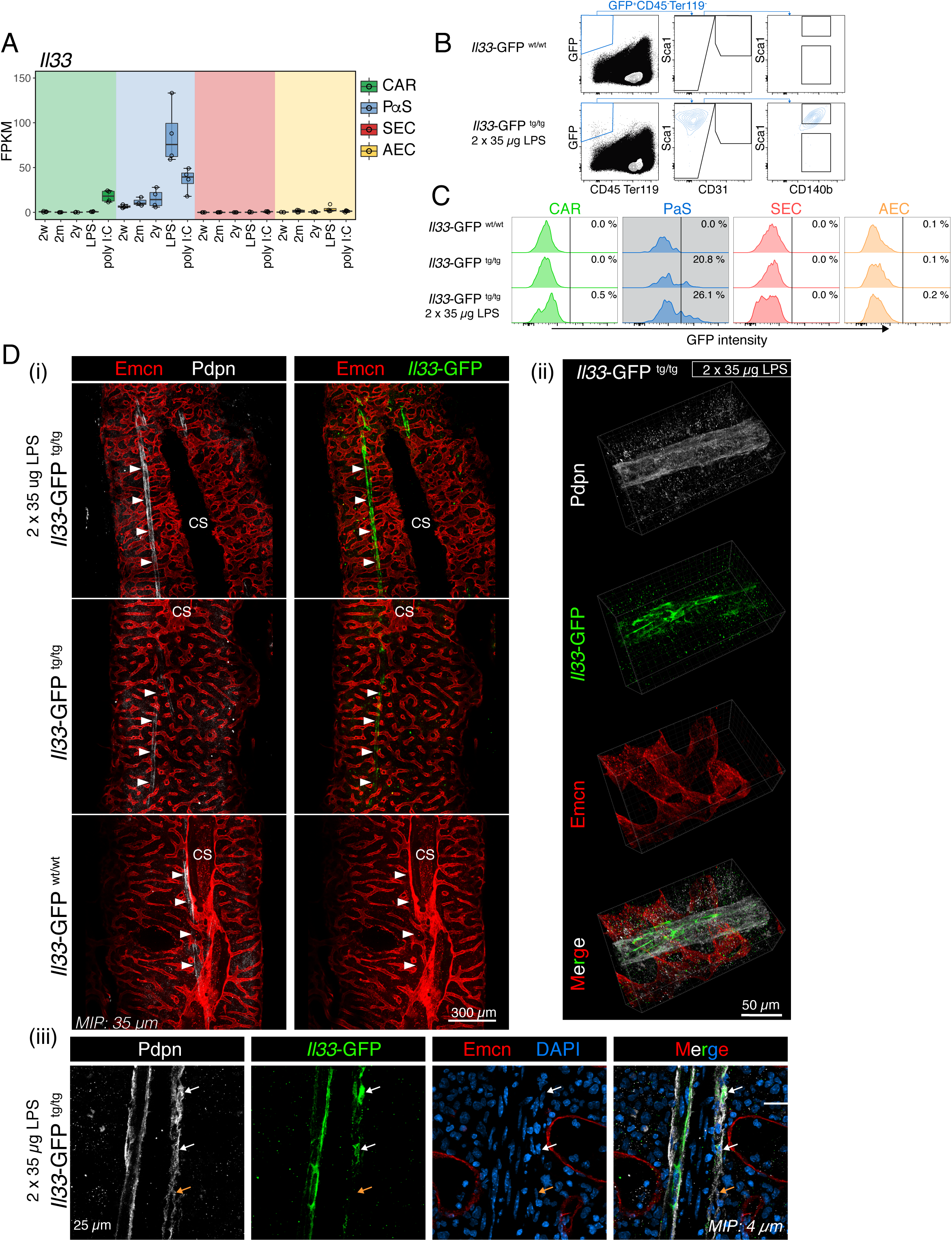
**A.** RNA-seq expression values (FPKM) for *Il33*. **B and C**. Flow cytometry analysis of *Il33*-GFP reporter mice. Pre-gated for “singlets” and “live cells” (DAPI^−^). **C**. Histograms displaying GFP intensity of pre-gated stromal cell populations. **D.** Immunofluorescence staining for Pdpn and Emcn on thick femoral slices of *Il33*-GFP reporter mice. (i) A diaphyseal region is shown, highlighting centrally located arteries (white arrow heads). cs: Central sinus. Scale bar: 300 *µ*m. (ii) Volumetric high-resolution rendering of a diaphyseal region containing a large artery. Scale bar: 50 *µ*m. (iii) High resolution optical 4 *µ*m section displaying peri-arterial Pdpn^+^ *Il33*-GFP^+^ cells. Scale bar: 25 *µ*m. MIP: Maximum intensity projection of indicated thickness

**Figure S4. Related to Figure 3.**
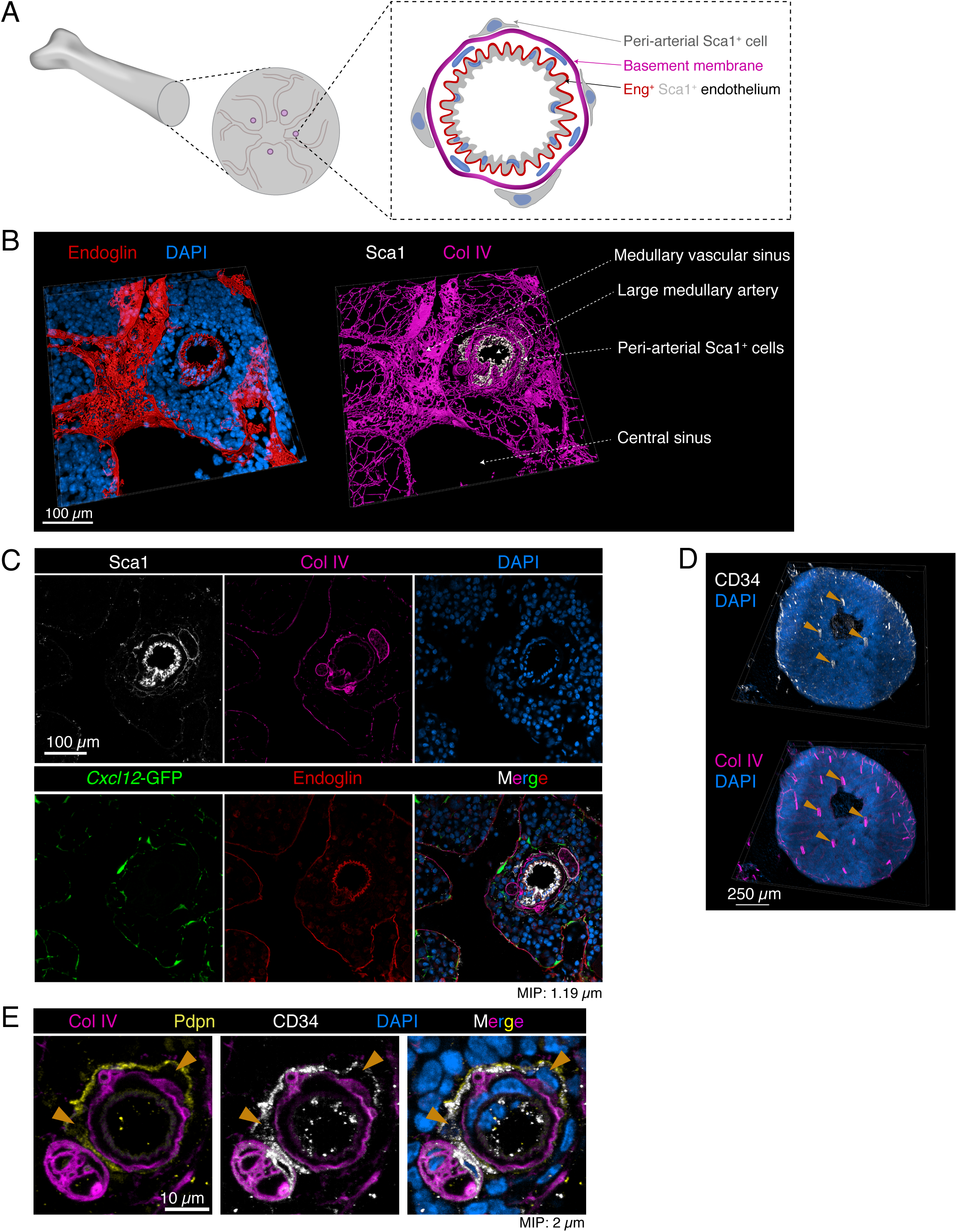
**A.** Scheme displaying the radial location of large arteries around the central sinus. Large arteries display a thick Collagen IV^+^ basement membrane ensheathed by an adventitial Sca1^+^ cell layer. **B.** Volumetric rendering of an immunostained transversal BM section. Endomucin and Collagen IV and Sca1 are displayed as segmented surfaces for better visualization. Scale bar: 100 *µ*m. **C.** Thin optical section (1.19 *µ*m) of the image in B. Scale bar: 100 *µ*m. MIP: Maximum intensity projection. **D.** Volumetric rendering of an immunostained thick transversal femoral slice. Scale bar: 250 *µ*m. **E.** High magnification image of a thin optical section (2 *µ*m) through a large BM artery. Pdpn^+^ CD34^+^ cells (arrow heads) are located outside of the Col IV^+^ basement membrane. Scale bar: 10 *µ*m.

**Figure S5. Related to Figure 5.**
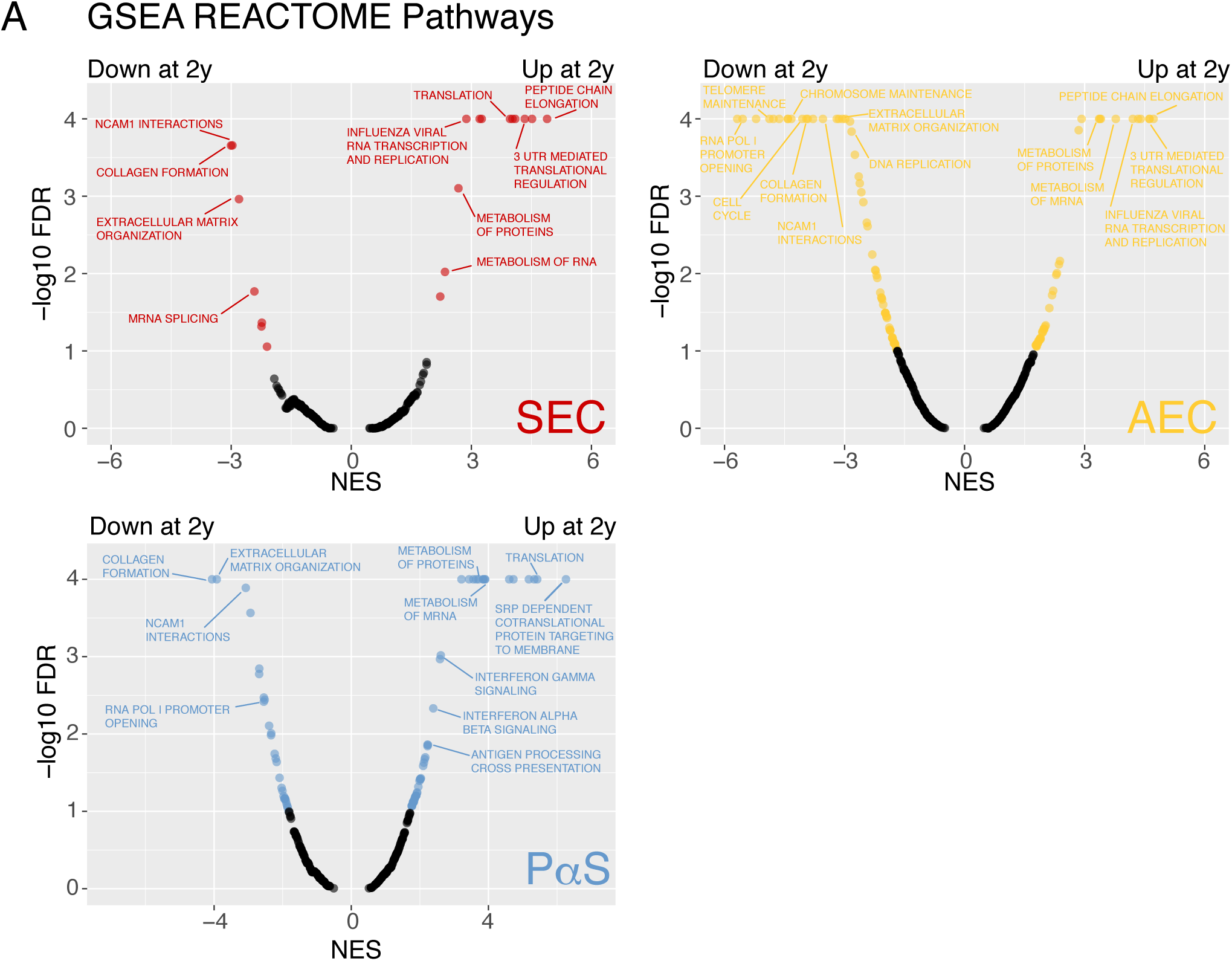
**A**. GSEA results for all REACTOME pathways comparing aged to adult stromal cells. Significant gene sets are displayed in color (FDR ≤ 0.05). NES: Normalized enrichment score

**Figure S6. Related to Figure 6.**
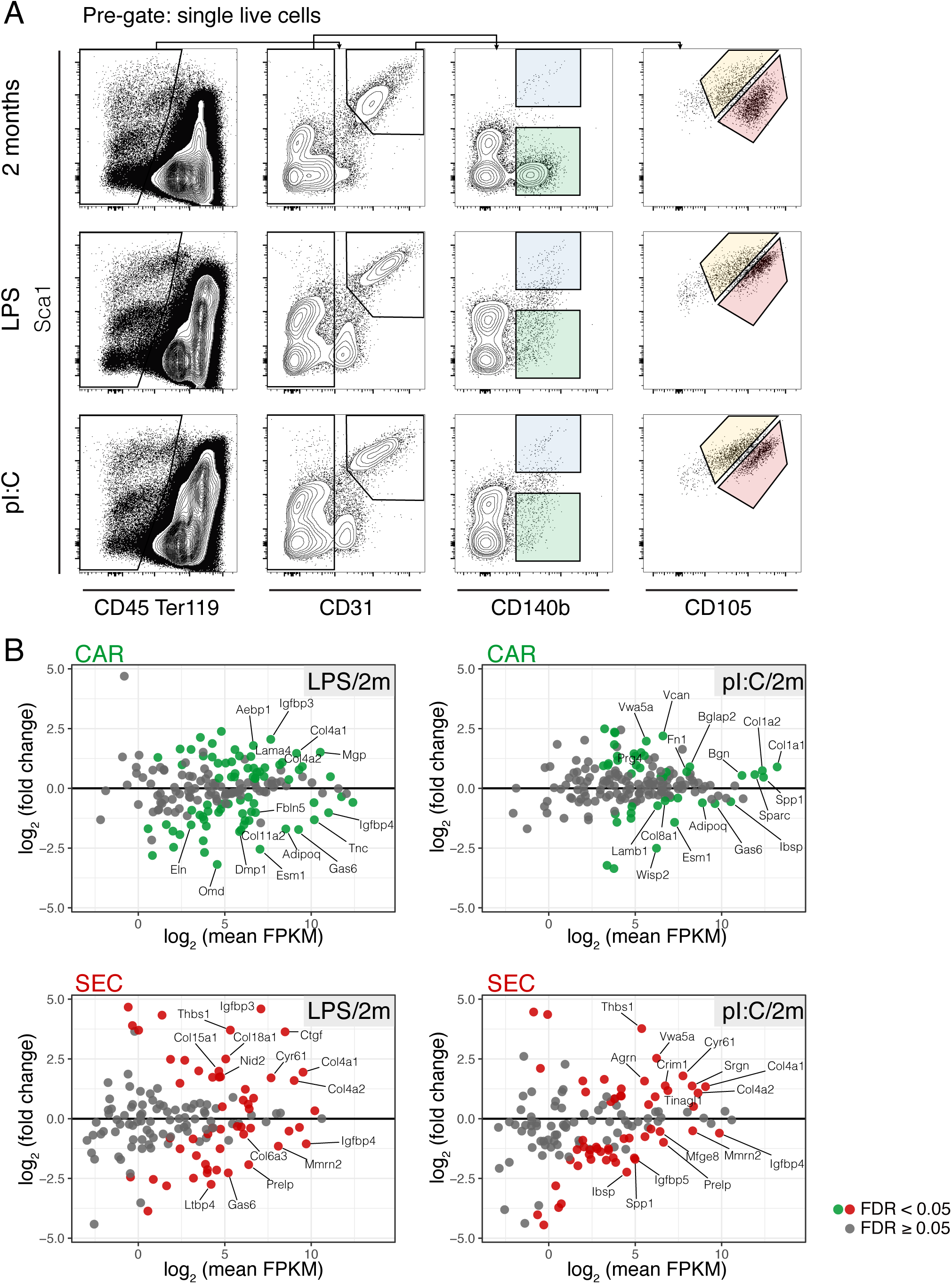
**A.** FACS gating strategy of bone marrow stromal cell populations for the conditions 2 weeks, LPS, and pI:C. Pre-gated for “singlets” and “live cells” (DAPI^−^). **B.** All detected core matrisome genes for CARc and SECs are displayed. The Y-axis shows the log2 FC of the pairwise comparisons LPS vs 2m or pI:C vs 2m. Significantly DEGs (FDR ≤ 0.05) are highlighted in color.

## References

Acar, M., Kocherlakota, K.S., Murphy, M.M., Peyer, J.G., Oguro, H., Inra, C.N., Jaiyeola, C., Zhao, Z., Luby-Phelps, K., Morrison, S.J., 2015. Deep imaging of bone marrow shows non-dividing stem cells are mainly perisinusoidal. Nature 526, 126–130. doi:10.1038/nature15250

Adkins, B., Leclerc, C., Marshall-Clarke, S., 2004. Neonatal adaptive immunity comes of age. Nature Reviews Immunology 4, 553–564. doi:10.1038/nri1394

Anders, S., Pyl, P.T., Huber, W., 2015. HTSeq--a Python framework to work with high-throughput sequencing data. Bioinformatics 31, 166–169. doi:10.1093/bioinformatics/btu638

Andrews, S., 2010. FastQC: a quality control tool for high throughput sequence data. Babraham Institute, Cambridge, United Kingdom.

Ara, T., Tokoyoda, K., Sugiyama, T., Egawa, T., Kawabata, K., Nagasawa, T., 2003. Long-Term Hematopoietic Stem Cells Require Stromal Cell-Derived Factor-1 for Colonizing Bone Marrow during Ontogeny. Immunity 19, 257–267. doi:10.1016/S1074-7613(03)00201-2

Arora, N., Wenzel, P.L., McKinney-Freeman, S.L., Ross, S.J., Kim, P.G., Chou, S.S., Yoshimoto, M., Yoder, M.C., Daley, G.Q., 2014. Effect of Developmental Stage of HSC and Recipient on Transplant Outcomes. Developmental Cell 29, 621–628. doi:10.1016/j.devcel.2014.04.013

Baryawno, N., Przybylski, D., Kowalczyk, M.S., Kfoury, Y., Severe, N., Gustafsson, K., Kokkaliaris, K.D., Mercier, F., Tabaka, M., Hofree, M., Dionne, D., Papazian, A., Lee, D., Ashenberg, O., Subramanian, A., Vaishnav, E.D., Rozenblatt-Rosen, O., Regev, A., Scadden, D.T., 2019. A Cellular Taxonomy of the Bone Marrow Stroma in Homeostasis and Leukemia. Cell 1–35. doi:10.1016/j.cell.2019.04.040

Beerman, I., Maloney, W.J., Weissmann, I.L., Rossi, D.J., 2010. Stem cells and the aging hematopoietic system. Current Opinion in Immunology 22, 500–506. doi:10.1016/j.coi.2010.06.007

Benz, C., Copley, M.R., Kent, D.G., Wohrer, S., Cortes, A., Aghaeepour, N., Ma, E., Mader, H., Rowe, K., Day, C., Treloar, D., Brinkman, R.R., Eaves, C.J., 2012. Hematopoietic stem cell subtypes expand differentially during development and display distinct lymphopoietic programs. Cell Stem Cell 10, 273– 283. doi:10.1016/j.stem.2012.02.007

Berg, J.S., Lin, K.K., Sonnet, C., Boles, N.C., Weksberg, D.C., Nguyen, H., Holt, L.J., Rickwood, D., Daly, R.J., Goodell, M.A., 2011. Imprinted Genes That Regulate Early Mammalian Growth Are Coexpressed in Somatic Stem Cells. PLoS ONE 6, e26410–10. doi:10.1371/journal.pone.0026410

Bernitz, J.M., Kim, H.S., MacArthur, B., Sieburg, H., Moore, K., 2016. Hematopoietic stem cells count and remember self-renewal divisions. Cell. doi:10.1016/j.cell.2016.10.022

Bixel, M.G., Kusumbe, A.P., Ramasamy, S.K., Sivaraj, K.K., Butz, S., Vestweber, D., Adams, R.H., 2017. Flow Dynamics and HSPC Homing in Bone Marrow Microvessels. CellReports 18, 1804–1816. doi:10.1016/j.celrep.2017.01.042

Boeuf, S., Kunz, P., Hennig, T., Lehner, B., Hogendoorn, P., Bovée, J., Richter, W., 2008. A chondrogenic gene expression signature in mesenchymal stem cells is a classifier of conventional central chondrosarcoma. J. Pathol. 216, 158–166. doi:10.1002/path.2389

Bowie, M.B., McKnight, K.D., Kent, D.G., McCaffrey, L., Hoodless, P.A., Eaves, C.J., 2006. Hematopoietic stem cells proliferate until after birth and show a reversible phase-specific engraftment defect. Journal of Clinical Investigation 116, 2808–2816. doi:10.1172/JCI28310DS1

Butler, J.M., Nolan, D.J., Vertes, E.L., Varnum-Finney, B., Kobayashi, H., Hooper, A.T., Seandel, M., Shido, K., White, I.A., Kobayashi, M., Witte, L., May, C., Shawber, C., Kimura, Y., Kitajewski, J., Rosenwaks, Z., Bernstein, I.D., Rafii, S., 2010. Endothelial cells are essential for the self-renewal and repopulation of Notch-dependent hematopoietic stem cells. Cell Stem Cell 6, 251–264. doi:10.1016/j.stem.2010.02.001

Casanova-Acebes, M., Pitaval, C., Weiss, L.A., Nombela Arrieta, C., Chèvre, R., A-González, N., Kunisaki, Y., Zhang, D., van Rooijen, N., Silberstein, L.E., Weber, C., Nagasawa, T., Frenette, P.S., Castrillo, A., Hidalgo, A., 2013. Rhythmic Modulation of the Hematopoietic Niche through Neutrophil Clearance. Cell 153, 1025–1035. doi:10.1016/j.cell.2013.04.040

Chambers, S.M., Boles, N.C., Lin, K.-Y.K., Tierney, M.P., Bowman, T.V., Bradfute, S.B., Chen, A.J., Merchant, A.A., Sirin, O., Weksberg, D.C., Merchant, M.G., Fisk, C.J., Shaw, C.A., Goodell, M.A., 2007a. Hematopoietic Fingerprints: An Expression Database of Stem Cells and Their Progeny. Cell Stem Cell 1, 578–591. doi:10.1016/j.stem.2007.10.003

Chambers, S.M., Shaw, C.A., Gatza, C., Fisk, C.J., Donehower, L.A., Goodell, M.A., 2007b. Aging Hematopoietic Stem Cells Decline in Function and Exhibit Epigenetic Dysregulation. Plos Biol 5, e201– 13. doi:10.1371/journal.pbio.0050201

Chavakis, T., Mitroulis, I., Hajishengallis, G., 2019. Hematopoietic progenitor cells as integrative hubs for adaptation to and fine-tuning of inflammation. Nature Immunology 20, 1–10. doi:10.1038/s41590-019-0402-5

Choi, J.S., Harley, B.A.C., 2017. Marrow-inspired matrix cues rapidly affect early fate decisions of hematopoietic stem and progenitor cells. Sci Adv 3, e1600455. doi:10.1126/sciadv.1600455

Chou, S., Lodish, H.F., 2010. Fetal liver hepatic progenitors are supportive stromal cells for hematopoietic stem cells. Proceedings of the National Academy of Sciences 107, 7799–7804. doi:10.1073/pnas.1003586107

Chow, A., Lucas, D., Hidalgo, A., Mendez-Ferrer, S., Hashimoto, D., Scheiermann, C., Battista, M., Leboeuf, M., Prophete, C., van Rooijen, N., Tanaka, M., Merad, M., Frenette, P.S., 2011. Bone marrow CD169+ macrophages promote the retention of hematopoietic stem and progenitor cells in the mesenchymal stem cell niche. Journal of Experimental Medicine 208, 261–271. doi:10.1084/jem.20101688

Corada, M., Orsenigo, F., Morini, M.F., Pitulescu, M.E., Bhat, G., Nyqvist, D., Breviario, F., Conti, V., Briot, A., Iruela-Arispe, M.L., Adams, R.H., Dejana, E., 1AD. Sox17 is indispensable for acquisition and maintenance of arterial identity. Nature Communications 4, 1–14. doi:10.1038/ncomms3609

Dahlgren, M.W., Jones, S.W., Cautivo, K.M., Dubinin, A., Ortiz-Carpena, J.F., Farhat, S., Yu, K.S., Lee, K., Wang, C., Molofsky, A.V., Tward, A.D., Krummel, M.F., Peng, T., Molofsky, A.B., 2019. Adventitial Stromal Cells Define Group 2 Innate Lymphoid Cell Tissue Niches. Immunity 50, 707–722.e6. doi:10.1016/j.immuni.2019.02.002

de Haan, G., Lazare, S.S., 2018. Aging of hematopoietic stem cells. Blood 131, 479–487. doi:10.1182/blood-2017-06-746412

de Magalhães, J.P., Curado, J., Church, G.M., 2009. Meta-analysis of age-related gene expression profiles identifies common signatures of aging. Bioinformatics 25, 875–881. doi:10.1093/bioinformatics/btp073

Ding, L., Saunders, T.L., Enikolopov, G., Morrison, S.J., 2012. Endothelial and perivascular cells maintain haematopoietic stem cells. Nature 481, 457–462. doi:10.1038/nature10783

Eaves, C.J., 2015. Hematopoietic stem cells: concepts, definitions, and the new reality. Blood 125, 2605–2613. doi:10.1182/blood-2014-12-570200

Ernst, J., bioinformatics, Z.B.-J.B., 2006, n.d. STEM: a tool for the analysis of short time series gene expression data. bmcbioinformatics. biomedcentral …. doi:10.1186/1471-2105-7-191

Espin-Palazon, R., Weijts, B., Mulero, V., Traver, D., 2018. Proinflammatory Signals as Fuel for the Fire of Hematopoietic Stem Cell Emergence. Trends in Cell Biology 28, 58–66. doi:10.1016/j.tcb.2017.08.003

Essers, M.A.G., Offner, S., Blanco-Bose, W.E., Waibler, Z., Kalinke, U., Duchosal, M.A., Trumpp, A., 2009. IFNα activates dormant haematopoietic stem cells in vivo. Nature 1–6. doi:10.1038/nature07815

Gazit, R., Garrison, B.S., Rao, T.N., Shay, T., Costello, J., Ericson, J., Kim, F., Collins, J.J., Regev, A., Wagers, A.J., Rossi, D.J., Consortium, T.I.G.P., 2013. Transcriptome Analysis Identifies Regulators of Hematopoietic Stem and Progenitor Cells. Stem Cell Reports 1, 266–280. doi:10.1016/j.stemcr.2013.07.004

Geiger, H., de Haan, G., Florian, M.C., 2013. The ageing haematopoietic stem cell compartment. Nature Reviews Immunology 13, 376–389. doi:10.1038/nri3433

Gomariz, Á., Helbling, P.M., Isringhausen, S., Suessbier, U., Becker, A., Boss, A., Nagasawa, T., Paul, G., Goksel, O., Székely, G., Stoma, S., Nørrelykke, S.F., Manz, M.G., Nombela Arrieta, C., 2018. Quantitative spatial analysis of haematopoiesis-regulating stromal cells in the bone marrow microenvironment by 3D microscopy. Nature Communications 9, 407–15. doi:10.1038/s41467-018-04770-z

Gomes, A.C., Gomes, M.S., 2016. Hematopoietic niches, erythropoiesis and anemia of chronic infection 1–7. doi:10.1016/j.exphem.2015.11.007

Graña, O., Rubio-Camarillo, M., Fdez-Riverola, F., Pisano, D.G., Glez-Peña, D., 2018. Nextpresso: Next generation sequencing expression analysis pipeline. Current Bioinformatics 13, 583–591. doi:10.2174/1574893612666170810153850

Greenbaum, A., Hsu, Y.-M.S., Day, R.B., Schuettpelz, L.G., Christopher, M.J., Borgerding, J.N., Nagasawa, T., Link, D.C., 2013. CXCL12 in early mesenchymal progenitors is required for haematopoietic stem-cell maintenance. Nature 495, 227–230. doi:10.1038/nature11926

He, Q., Zhang, C., Wang, L., Zhang, P., Ma, D., Lv, J., Liu, F., 2015. Inflammatory signaling regulates hematopoietic stem and progenitor cell emergence in vertebrates. Blood 125, 1098–1106. doi:10.1182/blood-2014-09-601542

Henry, C.J., Casás-Selves, M., Kim, J., Zaberezhnyy, V., Aghili, L., Daniel, A.E., Jimenez, L., Azam, T., McNamee, E.N., Clambey, E.T., Klawitter, J., Serkova, N.J., Tan, A.C., Dinarello, C.A., DeGregori, J., 2015. Aging-associated inflammation promotes selection for adaptive oncogenic events in B cell progenitors. J. Clin. Invest. doi:10.1172/JCI83024DS1

Himburg, H.A., Termini, C.M., Schlussel, L., Kan, J., Li, M., Zhao, L., Fang, T., Sasine, J.P., Chang, V.Y., Chute, J.P., 2018. Distinct Bone Marrow Sources of Pleiotrophin Control Hematopoietic Stem Cell Maintenance and Regeneration. Cell Stem Cell 23, 370–381.e5. doi:10.1016/j.stem.2018.07.003

Hu, X., Garcia, M., Weng, L., Jung, X., Murakami, J.L., Kumar, B., Warden, C.D., Todorov, I., Chen, C.-C., 2016. Identification of a common mesenchymal stromal progenitor for the adult haematopoietic niche. Nature Communications 7, 13095. doi:10.1038/ncomms13095

Itkin, T., Gur-Cohen, S., Spencer, J.A., Schajnovitz, A., Ramasamy, S.K., Kusumbe, A.P., Ledergor, G., Jung, Y., Milo, I., Poulos, M.G., Kalinkovich, A., Ludin, A., Kollet, O., Shakhar, G., Butler, J.M., Rafii, S., Adams, R.H., Scadden, D.T., Lin, C.P., Lapidot, T., 2016. Distinct bone marrow blood vessels differentially regulate haematopoiesis. Nature 532, 323–328. doi:10.1038/nature17624

Kfoury, Y., Scadden, D.T., 2015. Mesenchymal cell contributions to the stem cell niche. Cell Stem Cell 16, 239–253. doi:10.1016/j.stem.2015.02.019

Ki, S., Park, D., Selden, H.J., Seita, J., Chung, H., Kim, J., Iyer, V.R., Ehrlich, L.I.R., 2014. Global transcriptional profiling reveals distinct functions of thymic stromal subsets and age-related changes during thymic involution. CellReports 9, 402–415. doi:10.1016/j.celrep.2014.08.070

Kim, I., Saunders, T.L., Morrison, S.J., 2007. Sox17 Dependence Distinguishes the Transcriptional Regulation of Fetal from Adult Hematopoietic Stem Cells. Cell 130, 470–483. doi:10.1016/j.cell.2007.06.011

Kohara, H., Omatsu, Y., Sugiyama, T., Noda, M., Fujii, N., Nagasawa, T., 2007. Development of plasmacytoid dendritic cells in bone marrow stromal cell niches requires CXCL12-CXCR4 chemokine signaling. Blood 110, 4153–4160. doi:10.1182/blood-2007-04-084210

Kokkaliaris, K.D., Drew, E., Endele, M., Loeffler, D., Hoppe, P.S., Hilsenbeck, O., Schauberger, B., Hinzen, C., Skylaki, S., Theodorou, M., Kieslinger, M., Lemischka, I., Moore, K., Schroeder, T., 2016. Identification of factors promoting ex vivo maintenance of mouse hematopoietic stem cells by long-term single-cell quantification. Blood 128, 1181–1192. doi:10.1182/blood-2016-03-705590

Komuro, A.N.T.S.S.N.M.-L.L.T.H.A.N.K.O.T.S.A.H.Y.H.I.S.W.Z.H.T.A.K.H.A.T.O.J.-K.L.T.M.T.N.K.W.A.K.M.M.M.B.I.S.I., Sumida, T., Nomura, S., Liu, M.-L., Higo, T., Nakagawa, A., Okada, K., Sakai, T., Hashimoto, A., Hara, Y., Shimizu, I., Zhu, W., Toko, H., Katada, A., Akazawa, H., Oka, T., Lee, J.-K., Minamino, T., Nagai, T., Walsh, K., Kikuchi, A., Matsumoto, M., Botto, M., Shiojima, I., Komuro, I., 2012. Complement C1q Activates Canonical Wnt Signaling and Promotes Aging-Related Phenotypes. Cell 149, 1298–1313. doi:10.1016/j.cell.2012.03.047

Kovtonyuk, L.V., Fritsch, K., Feng, X., Manz, M.G., Takizawa, H., 2016. Inflamm-Aging of Hematopoiesis, Hematopoietic Stem Cells, and the Bone Marrow Microenvironment. Front Immunol 7, 405–13. doi:10.3389/fimmu.2016.00502

Kusumbe, A.P., Ramasamy, S.K., Adams, R.H., 2014. Coupling of angiogenesis and osteogenesis by a specific vessel subtype in bone. Nature 507, 323–328. doi:10.1038/nature13145

Kusumbe, A.P., Ramasamy, S.K., Itkin, T., Mäe, M.A., Langen, U.H., Betsholtz, C., Lapidot, T., Adams, R.H., 2016. Age-dependent modulation of vascular niches for haematopoietic stem cells. Nature 532, 380–384. doi:10.1038/nature17638

Langmead, B., Trapnell, C., Pop, M., Salzberg, S.L., 2009. Ultrafast and memory-efficient alignment of short DNA sequences to the human genome. Genome Biol 10, R25. doi:10.1186/gb-2009-10-3-r25

Lazzari, E., Butler, J.M., 2018. The Instructive Role of the Bone Marrow Niche in Aging and Leukemia 1–8. doi:10.1007/s40778-018-0143-7

Li, H., 2011. A statistical framework for SNP calling, mutation discovery, association mapping and population genetical parameter estimation from sequencing data. Bioinformatics 27, 2987–2993. doi:10.1093/bioinformatics/btr509

Li, X.M., Hu, Z., Jorgenson, M.L., Slayton, W.B., 2009. High Levels of Acetylated Low-Density Lipoprotein Uptake and Low Tyrosine Kinase With Immunoglobulin and Epidermal Growth Factor Homology Domains-2 (Tie2) Promoter Activity Distinguish Sinusoids From Other Vessel Types in Murine Bone Marrow. Circulation 120, 1910–1918. doi:10.1161/CIRCULATIONAHA.109.871574

Louis, I., Heinonen, K.M., Chagraoui, J., Vainio, S., Sauvageau, G., Perreault, C., 2008. The Signaling Protein Wnt4 Enhances Thymopoiesis and Expands Multipotent Hematopoietic Progenitors through β-Catenin-Independent Signaling. Immunity 29, 57–67. doi:10.1016/j.immuni.2008.04.023

Love, M.I., Huber, W., Anders, S., 2014. Moderated estimation of fold change and dispersion for RNA-seq data with DESeq2. Genome Biol 15, 31–21. doi:10.1186/s13059-014-0550-8

Mahlakõiv, T., Flamar, A.L., Johnston, L.K., Moriyama, S., Putzel, G.G., Bryce, P.J., Artis, D., 2019. Stromal cells maintain immune cell homeostasis in adipose tissue via production of interleukin-33. Sci Immunol 4, eaax0416. doi:10.1126/sciimmunol.aax0416

Manz, M.G., Boettcher, S., 2014. Emergency granulopoiesis. Nature Reviews Immunology 14, 302– 314. doi:10.1038/nri3660

Martin, M., 2011. Cutadapt removes adapter sequences from high-throughput sequencing reads. EMBnet.journal 17, 10–12. doi:10.14806/ej.17.1.200

Maryanovich, M., Zahalka, A.H., Pierce, H., Pinho, S., Nakahara, F., Asada, N., Wei, Q., Wang, X., Ciero, P., Xu, J., Leftin, A., Frenette, P.S., 2018. Adrenergic nerve degeneration in bone marrow drives aging of the hematopoietic stem cell niche. Nature Medicine 1–18. doi:10.1038/s41591-018-0030-x

Mascarenhas, M.I., Parker, A., Dzierzak, E., Ottersbach, K., 2009. Identification of novel regulators of hematopoietic stem cell development through refinement of stem cell localization and expression profiling. Blood 114, 4645–4653. doi:10.1182/blood-2009-06-230037

McGeer, E.G., Klegeris, A., McGeer, P.L., 2005. Inflammation, the complement system and the diseases of aging. Neurobiology of Aging 26, 94–97. doi:10.1016/j.neurobiolaging.2005.08.008

Mercier, F.E., Ragu, C., Scadden, D.T., 2012. The bone marrow at the crossroads of blood and immunity. Nature Reviews Immunology 12, 49–60. doi:10.1038/nri3132

Mi, H., Muruganujan, A., Ebert, D., Huang, X., Thomas, P.D., 2018. PANTHER version 14: more genomes, a new PANTHER GO-slim and improvements in enrichment analysis tools. Nucleic Acids Research 47, D419–D426. doi:10.1093/nar/gky1038

Mootha, V.K., Lindgren, C.M., Eriksson, K.-F., Subramanian, A., Sihag, S., Lehar, J., Puigserver, P., Carlsson, E., Ridderstråle, M., Laurila, E., Houstis, N., Daly, M.J., Patterson, N., Mesirov, J.P., Golub, T.R., Tamayo, P., Spiegelman, B., Lander, E.S., Hirschhorn, J.N., Altshuler, D., Groop, L.C., 2003. PGC-1alpha-responsive genes involved in oxidative phosphorylation are coordinately downregulated in human diabetes. Nature Genetics 34, 267–273. doi:10.1038/ng1180

Morikawa, S., Mabuchi, Y., Kubota, Y., Nagai, Y., Niibe, K., Hiratsu, E., Suzuki, S., Miyauchi-Hara, C., Nagoshi, N., Sunabori, T., Shimmura, S., Miyawaki, A., Nakagawa, T., Suda, T., Okano, H., Matsuzaki, Y., 2009. Prospective identification, isolation, and systemic transplantation of multipotent mesenchymal stem cells in murine bone marrow. Journal of Experimental Medicine 206, 2483–2496. doi:10.1084/jem.20091046

Moscatello, K.M., Biber, K.L., Dempsey, D.C., Chervenak, R., Wolcott, R.M., 1998. Characterization of a B cell progenitor present in neonatal bone marrow and spleen but not in adult bone marrow and spleen. J. Immunol. 161, 5391–5398.

Nakamura-Ishizu, A., Takizawa, H., Suda, T., 2014. The analysis, roles and regulation of quiescence in hematopoietic stem cells. Development 141, 4656–4666. doi:10.1242/dev.106575

Noda, M., Omatsu, Y., Sugiyama, T., Oishi, S., Fujii, N., Nagasawa, T., 2011. CXCL12-CXCR4 chemokine signaling is essential for NK-cell development in adult mice. Blood 117, 451–458. doi:10.1182/blood-2010-04-277897

Nombela Arrieta, C., Isringhausen, S., 2017. The Role of the Bone Marrow Stromal Compartment in the Hematopoietic Response to Microbial Infections. Front Immunol 7, 631–12. doi:10.3389/fimmu.2016.00689

Nombela-Arrieta, C., Manz, M.G., 2017. Quantification and three-dimensional microanatomical organization of the bone marrow. Blood Advances.

Nusspaumer, G., Jaiswal, S., Barbero, A., Reinhardt, R., Ronen, D.I., Haumer, A., Lufkin, T., Martin, I., Zeller, R., 2017. Ontogenic Identification and Analysis of Mesenchymal Stromal Cell Populations during Mouse Limb and Long Bone Development. Stem Cell Reports 1–33. doi:10.1016/j.stemcr.2017.08.007

Oboki, K., Ohno, T., Kajiwara, N., Arae, K., Morita, H., Ishii, A., Nambu, A., Abe, T., Kiyonari, H., Matsumoto, K., Sudo, K., Okumura, K., Saito, H., Nakae, S., 2010. IL-33 is a crucial amplifier of innate rather than acquired immunity. Proc. Natl. Acad. Sci. U.S.A. 107, 18581–18586. doi:10.1073/pnas.1003059107

Omatsu, Y., Seike, M., Sugiyama, T., Kume, T., Nagasawa, T., 2014. Foxc1 is a critical regulator of haematopoietic stem/ progenitor cell niche formation. Nature 1–16. doi:10.1038/nature13071

Omatsu, Y., Sugiyama, T., Kohara, H., Kondoh, G., Fujii, N., Kohno, K., Nagasawa, T., 2010. The Essential Functions of Adipo-osteogenic Progenitors as the Hematopoietic Stem and Progenitor Cell Niche. Immunity 33, 387–399. doi:10.1016/j.immuni.2010.08.017

Park, M.H., Jin, H.K., Min, W.K., Lee, W.W., Lee, J.E., Akiyama, H., Herzog, H., Enikolopov, G.N., Schuchman, E.H., Bae, J.S., 2015. Neuropeptide Y regulates the hematopoietic stem cell microenvironment and prevents nerve injury in the bone marrow. EMBO J. 34, 1648–1660. doi:10.15252/embj.201490174

Pietras, E.M., Mirantes-Barbeito, C., Fong, S., Loeffler, D., Kovtonyuk, L.V., Zhang, S., Lakshminarasimhan, R., Chin, C.P., Techner, J.-M., Will, B., Nerlov, C., Steidl, U., Manz, M.G., Schroeder, T., Passegué, E., 2016. Chronic interleukin-1 exposure drives haematopoietic stem cells towards precocious myeloid differentiation at the expense of self-renewal. Nat. Cell Biol. 18, 607–618. doi:10.1038/ncb3346

Pihlgren, M., Schallert, N., Tougne, C., Bozzotti, P., Kovarik, J., Fulurija, A., Kosco-Vilbois, M., Lambert, P.H., Siegrist, C.A., 2001. Delayed and deficient establishment of the long-term bone marrow plasma cell pool during early life. Eur. J. Immunol. 31, 939–946. doi:10.1002/1521-4141(200103)31:3<939::AID-IMMU939>3.0.CO;2-I

Ratajczak, M.Z., Kim, C.H., Wojakowski, W., Janowska-Wieczorek, A., Kucia, M., Ratajczak, J., 2010. Innate immunity as orchestrator of stem cell mobilization. Leukemia 24, 1667–1675. doi:10.1038/leu.2010.162

Salzer, M.C., Lafzi, A., Berenguer-Llergo, A., Youssif, C., Castellanos, A., Solanas, G., Peixoto, F.O., Attolini, C.S.-O., Prats, N., Aguilera, M., Martín-Caballero, J., Heyn, H., Benitah, S.A., 2018. Identity Noise and Adipogenic Traits Characterize Dermal Fibroblast Aging. Cell 175, 1575–1590.e22. doi:10.1016/j.cell.2018.10.012

Sawamiphak, S., Kontarakis, Z., Stainier, D.Y.R., 2014. Interferon Gamma Signaling Positively Regulates Hematopoietic Stem Cell Emergence. Developmental Cell 31, 640–653. doi:10.1016/j.devcel.2014.11.007

Schreck, C., Istvánffy, R., Ziegenhain, C., Sippenauer, T., Ruf, F., Henkel, L., Gärtner, F., Vieth, B., Florian, M.C., Mende, N., Taubenberger, A., Prendergast, Á., Wagner, A., Pagel, C., Grziwok, S., Götze, K.S., Guck, J., Dean, D.C., Massberg, S., Essers, M., Waskow, C., Geiger, H., Schiemann, M., Peschel, C., Enard, W., Oostendorp, R.A.J., 2017. Niche WNT5A regulates the actin cytoskeleton during regeneration of hematopoietic stem cells. J. Exp. Med. 214, 165–181. doi:10.1084/jem.20151414

Schumacher, A., Denecke, B., Braunschweig, T., Stahlschmidt, J., Ziegler, S., Brandenburg, L.-O., Stope, M.B., Martincuks, A., Vogt, M., Görtz, D., Camporeale, A., Poli, V., Müller-Newen, G., Brümmendorf, T.H., Ziegler, P., 2015. Angptl4 is upregulated under inflammatory conditions in the bone marrow of mice, expands myeloid progenitors, and accelerates reconstitution of platelets after myelosuppressive therapy. J Hematol Oncol 1–16. doi:10.1186/s13045-015-0152-2

Schürch, C.M., Riether, C., Ochsenbein, A.F., 2014. Cytotoxic CD8(+) T Cells Stimulate Hematopoietic Progenitors by Promoting Cytokine Release from Bone Marrow Mesenchymal Stromal Cells. Cell Stem Cell. doi:10.1016/j.stem.2014.01.002

Shi, C., Jia, T., Méndez-Ferrer, S., Hohl, T.M., Serbina, N.V., Lipuma, L., Leiner, I., Li, M.O., Frenette, P.S., Pamer, E.G., 2011. Bone Marrow Mesenchymal Stem and Progenitor Cells Induce Monocyte Emigration in Response to Circulating Toll-like Receptor Ligands. Immunity 34, 590–601. doi:10.1016/j.immuni.2011.02.016

Singh, P., Hoggatt, J., Kamocka, M.M., Mohammad, K.S., Saunders, M.R., Li, H., Speth, J., Carlesso, N., Guise, T.A., Pelus, L.M., 2017. Neuropeptide Y regulates a vascular gateway for hematopoietic stem and progenitor cells. Journal of Clinical Investigation 127, 4527–4540. doi:10.1172/JCI94687

Sitnik, K.M., Wendland, K., Weishaupt, H., Uronen-Hansson, H., White, A.J., Anderson, G., Kotarsky, K., Agace, W.W., 2016. Context-Dependent Development of Lymphoid Stroma from Adult CD34+ Adventitial Progenitors. CellReports 14, 2375–2388. doi:10.1016/j.celrep.2016.02.033

Sivaraj, K.K., Adams, R.H., 2016. Blood vessel formation and function in bone. Development 143, 2706–2715. doi:10.1242/dev.136861

Smith-Berdan, S., Nguyen, A., Hong, M.A., Forsberg, E.C., 2015. ROBO4-mediated vascular integrity regulates the directionality of hematopoietic stem cell trafficking. Stem Cell Reports 4, 255–268. doi:10.1016/j.stemcr.2014.12.013

Spallanzani, R.G., Zemmour, D., Xiao, T., Jayewickreme, T., Li, C., Bryce, P.J., Benoist, C., Mathis, D., 2019. Distinct immunocyte-promoting and adipocyte-generating stromal components coordinate adipose tissue immune and metabolic tenors. Sci Immunol 4, eaaw3658. doi:10.1126/sciimmunol.aaw3658

Stegner, D., vanEeuwijk, J.M.M., Angay, O., Gorelashvili, M.G., Semeniak, D., Pinnecker, J., Schmithausen, P., Meyer, I., Friedrich, M., Dütting, S., Brede, C., Beilhack, A., Schulze, H., Nieswandt, B., Heinze, K.G., 2017. Thrombopoiesis is spatially regulated by the bone marrow vasculature. Nature Communications 1–11. doi:10.1038/s41467-017-00201-7

Subramanian, A., Tamayo, P., Mootha, V.K., Mukherjee, S., Ebert, B.L., Gillette, M.A., Paulovich, A., Pomeroy, S.L., Golub, T.R., Lander, E.S., Mesirov, J.P., 2005. Gene set enrichment analysis: a knowledge-based approach for interpreting genome-wide expression profiles. Proceedings of the National Academy of Sciences 102, 15545–15550. doi:10.1073/pnas.0506580102

Sugiyama, T., Kohara, H., Noda, M., Nagasawa, T., 2006. Maintenance of the Hematopoietic Stem Cell Pool by CXCL12-CXCR4 Chemokine Signaling in Bone Marrow Stromal Cell Niches. Immunity 25, 977–988. doi:10.1016/j.immuni.2006.10.016

Sun, D., Luo, M., Jeong, M., Rodriguez, B., Xia, Z., Hannah, R., Wang, H., Le, T., Faull, K.F., Chen, R., Gu, H., Bock, C., Meissner, A., Göttgens, B., Darlington, G.J., Li, W., Goodell, M.A., 2014. Epigenomic profiling of young and aged HSCs reveals concerted changes during aging that reinforce self-renewal. Cell Stem Cell 14, 673–688. doi:10.1016/j.stem.2014.03.002

Takizawa, H., Boettcher, S., Manz, M.G., 2012. Demand-adapted regulation of early hematopoiesis in infection and inflammation. Blood 119, 2991–3002. doi:10.1182/blood-2011-12-380113

Takizawa, H., Fritsch, K., Kovtonyuk, L.V., Saito, Y., Yakkala, C., Jacobs, K., Ahuja, A.K., Lopes, M., Hausmann, A., Hardt, W.-D., Gomariz, Á., Nombela Arrieta, C., Manz, M.G., 2017. Pathogen-Induced TLR4-TRIF Innate Immune Signaling in Hematopoietic Stem Cells Promotes Proliferation but Reduces Competitive Fitness. Cell Stem Cell 21, 225–240.e5. doi:10.1016/j.stem.2017.06.013

Takizawa, H., Regoes, R.R., Boddupalli, C.S., Bonhoeffer, S., Manz, M.G., 2011. Dynamic variation in cycling of hematopoietic stem cells in steady state and inflammation. Journal of Experimental Medicine 208, 273–284. doi:10.1084/jem.20101643

Tanegashima, K., Suzuki, K., Nakayama, Y., Tsuji, K., Shigenaga, A., Otaka, A., Hara, T., 2013. CXCL14 is a natural inhibitor of the CXCL12-CXCR4 signaling axis. FEBS Letters 587, 1731–1735. doi:10.1016/j.febslet.2013.04.046

Thomas, D.D., Sommer, A.G., Balazs, A.B., Beerman, I., Murphy, G.J., Rossi, D., Mostoslavsky, G., 2016. Insulin-like growth factor 2 modulates murine hematopoietic stem cell maintenance through upregulation of p57. Experimental Hematology 44, 422–433.e1. doi:10.1016/j.exphem.2016.01.010

Tikhonova, A.N., Dolgalev, I., Hu, H., Sivaraj, K.K., Hoxha, E., nguez, A.X.L.C.-D.X., Pinho, S., Akhmetzyanova, I., Gao, J., Witkowski, M., Guillamot, M., Gutkin, M.C., Zhang, Y., Marier, C., Diefenbach, C., Kousteni, S., Heguy, A., Zhong, H., Fooksman, D.R., Butler, J.M., Economides, A., Frenette, P.S., Adams, R.H., Satija, R., Tsirigos, A., Aifantis, I., 2019. The bone marrow microenvironment at single-cell resolution. Nature 505, 1–28. doi:10.1038/s41586-019-1104-8

Tokoyoda, K., Egawa, T., Sugiyama, T., Choi, B.-I., Nagasawa, T., 2004. Cellular Niches Controlling B Lymphocyte Behavior within Bone Marrow during Development. Immunity 20, 707–718. doi:10.1016/j.immuni.2004.05.001

Trapnell, C., Roberts, A., Goff, L., Pertea, G., Kim, D., Kelley, D.R., Pimentel, H., Salzberg, S.L., Rinn, J.L., Pachter, L., 2012. Differential gene and transcript expression analysis of RNA-seq experiments with TopHat and Cufflinks. Nat Protoc 7, 562–578. doi:10.1038/nprot.2012.016

Trompouki, E., 2016. Stress and Non-Stress Roles of Inflammatory Signals during HSC Emergence and Maintenance 1–15. doi:10.3389/fimmu.2016.00487&domain=pdf&date_stamp=2016-11-07

Venkatraman, A., He, X.C., Thorvaldsen, J.L., Sugimura, R., Perry, J.M., Tao, F., Zhao, M., Christenson, M.K., Sanchez, R., Yu, J.Y., Peng, L., Haug, J.S., Paulson, A., Li, H., Zhong, X.-B., Clemens, T.L., Bartolomei, M.S., Li, L., 2013. Maternal imprinting at the H19-Igf2 locus maintains adult haematopoietic stem cell quiescence. Nature 500, 345–349. doi:10.1038/nature12303

Walter, D., Lier, A., Geiselhart, A., Thalheimer, F.B., Huntscha, S., Sobotta, M.C., Moehrle, B., Brocks, D., Bayindir, I., Kaschutnig, P., Muedder, K., Klein, C., Jauch, A., Schroeder, T., Geiger, H., Dick, T.P., Holland-Letz, T., Schmezer, P., Lane, S.W., Rieger, M.A., Essers, M.A.G., Williams, D.A., Trumpp, A., Milsom, M.D., 2015. Exit from dormancy provokes DNA-damage-induced attrition in haematopoietic stem cells. Nature 520, 549–552. doi:10.1038/nature14131

Wilson, A., Laurenti, E., Oser, G., van der Wath, R.C., Blanco-Bose, W., Jaworski, M., Offner, S., Dunant, C.F., Eshkind, L., Bockamp, E., Pietro Lió, MacDonald, H.R., Trumpp, A., 2008. Hematopoietic Stem Cells Reversibly Switch from Dormancy to Self-Renewal during Homeostasis and Repair. Cell 135, 1118–1129. doi:10.1016/j.cell.2008.10.048

Wingett, S.W., Andrews, S., 2018. FastQ Screen: A tool for multi-genome mapping and quality control. F1000Res 7, 1338–13. doi:10.12688/f1000research.15931.1

Wu, Q., Kawahara, M., Kono, T., 2008. Synergistic role of Igf2 and Dlk1 in fetal liver development and hematopoiesis in bi-maternal mice. J. Reprod. Dev. 54, 177–182. doi:10.1262/jrd.19146

Xiao, Y., 2015. Loss of Angiopoietin-like 7 diminishes the regeneration capacity of hematopoietic stem and progenitor cells 1–5. doi:10.1186/s13045-014-0102-4

Xiao, Y., Jiang, Z., Li, Y., Ye, W., Jia, B., Zhang, M., Xu, Y., Wu, D., Lai, L., Chen, Y., Chang, Y., Huang, X., Liu, H., Qing, G., Liu, P., Li, Y., Li, Y., Xu, B., Zhong, M., Yao, Y., Pei, D., Li, P., 2015. ANGPTL7 regulates the expansion and repopulation of human hematopoietic stem and progenitor cells. Haematologica 100, 585–594. doi:10.3324/haematol.2014.118612

Xu, C., Gao, X., Wei, Q., Nakahara, F., Zimmerman, S.E., Mar, J., Frenette, P.S., 2018. Stem cell factor is selectively secreted by arterial endothelial cells in bone marrow. Nature Communications 1– 13. doi:10.1038/s41467-018-04726-3

Ye, M., Zhang, H., Amabile, G., Yang, H., Staber, P.B., Zhang, P., Levantini, E., Alberich-Jordà, M., Zhang, J., Kawasaki, A., Tenen, D.G., 2013. C/EBPa controls acquisition and maintenance of adult haematopoietic stem cell quiescence. Nat. Cell Biol. 15, 385–394. doi:10.1038/ncb2698

Zhang, H., Rodriguez, S., Wang, L., Wang, S., Serezani, H., Kapur, R., Cardoso, A.A., Carlesso, N., 2016. Sepsis Induces Hematopoietic Stem Cell Exhaustion and Myelosuppression through Distinct Contributions of TRIF and MYD88. Stem Cell Reports 6, 940–956. doi:10.1016/j.stemcr.2016.05.002

Zhou, B.O., Yue, R., Murphy, M.M., Peyer, J.G., Morrison, S.J., 2014. Leptin-receptor-expressing mesenchymal stromal cells represent the main source of bone formed by adult bone marrow. Cell Stem Cell 15, 154–168. doi:10.1016/j.stem.2014.06.008

