## Supplementary Table 3 for "Global transcriptomic profiling of the bone marrow stromal microenvironment during postnatal development, aging and inflammation"

**Table S2 – Materials**

| Name | Company | Identifier | Description |
| --- | --- | --- | --- |
| Histology antibodies |  |  |  |
| Podoplanin Antibody (eBio8.1.1) | ThermoFisher | 14-5381-85 | Primary ab. [5 µg/mL] |
| F4/80 Monoclonal Antibody (BM8) | ThermoFisher | 14-4801-82 | Primary ab. [5 µg/mL] |
| Anti-Collagen IV antibody | Abcam | ab6586 | Primary ab. [2.5 µg/mL] |
| Endomucin Antibody (V.7C7) | Santa Cruz Biotech. | Sc-65495 | Primary ab. [2 µg/mL] |
| Purified anti-mouse Ly-6A/E (Sca-1) Ab | BioLegend | 122502 | Primary ab. [5 µg/mL] |
| CD34 Monoclonal Antibody (RAM34) | ThermoFisher | 14-0341-81 | Primary ab. [5 µg/mL] |
| CD140b (PDGFRB), APC | ThermoFisher | 17-1402-82 | Primary ab. [4 µg/mL] |
| Biotin Goat Anti-Syrian Hamster IgG | Jackson Immuno R. | 107-065-142 | Secondary ab. [6 µg/mL] |
| Donkey anti-Goat IgG (H+L), AF546 | ThermoFisher | A-11056 | Secondary ab. [20 µg/mL] |
| Donkey anti-Goat IgG (H+L), AF680 | ThermoFisher | A-21084 | Secondary ab. [20 µg/mL] |
| Donkey anti-Rat IgG (H+L), AF594 | ThermoFisher | A-21209 | Secondary ab. [20 µg/mL] |
| Donkey anti-Rat IgG (H+L), AF488 | ThermoFisher | A-21208 | Secondary ab. [20 µg/mL] |
| Donkey anti-Rabbit IgG (H+L), AF680 | ThermoFisher | A10043 | Secondary ab. [20 µg/mL] |
| Donkey anti-Rabbit IgG (H+L), AF546 | ThermoFisher | A10040 | Secondary ab. [20 µg/mL] |
| Streptavidin, Alexa Fluor™ 555 conjugate | ThermoFisher | S32355 | Histology [20 µg/mL] |
| Flow cytometry antibodies |  |  |  |
| TruStain fcX™ (anti-mouse CD16/32) | BioLegend | 101320 | Flow cytometry [2.5 µg/mL] |
| CD45 (30-F11), PerCP-Cyanine5.5 | ThermoFisher | 45-0451-82 | Flow cytometry [1 µg/mL] |
| TER-119, PerCP-Cyanine5.5 | ThermoFisher | 45-5921-82 | Flow cytometry [1 µg/mL] |
| CD31 (PECAM-1), PE-Cyanine7 | ThermoFisher | 25-0311-82 | Flow cytometry [0.67 µg/mL] |
| CD140b (PDGFRB), APC | ThermoFisher | 17-1402-82 | Flow cytometry [4 µg/mL] |
| BV786 Rat Anti-Mouse CD105 (MJ7/18) | BD Biosciences | 564746 | Flow cytometry [1 µg/mL] |
| APC/Cy7 anti-mouse Ly-6A/E (Sca-1) | BioLegend | 108126 | Flow cytometry [0.4 µg/mL] |
| BV711 anti-mouse CD206 (MMR) Antibody | BioLegend | 141727 | Flow cytometry [2 µg/mL] |
| BV 711 Rat IgG2a, κ Isotype Ctrl Antibody | BioLegend | 400551 | Flow cytometry [2 µg/mL] |
| Podoplanin-PE Antibody (eBio8.1.1) | ThermoFisher | 12-5381-82 | Flow cytometry [2 µg/mL] |
| Syrian Hamster IgG-PE Isotype Control | ThermoFisher | 12-4914-81 | Flow cytometry [2 µg/mL] |
| PE Rat anti-Mouse CD34 (RAM34) | BD Biosciences | 551387 | Flow cytometry [2 µg/mL] |
| Chemicals & enzymes & consumables |  |  |  |
| LPS-EB Ultrapure | InvivoGen | tlrl-3pelps | Ultrap. LPS, E. coli 0111:B4 |
| Poly(I:C) (HMW) | InvivoGen | tlrl-pic-5 | Synth. analog of dsRNA |
| RapiClear® 1.52 | SunJin Lab | RC152001 | Tissue clearing |
| Collagenase, Type 2 | Worthington Bio. | LS004176 | Enzymatic tissue digestion |
| Deoxyribonuclease I | Worthington Bio. | LS002007 | Enzymatic tissue digestion |
| EDTA solution pH 8.0 (0.5 M) | Panreac AppliChem | A4892,0100 | Flow cytometry buffer |
| DMEM, high glucose, GlutaMAX suppl. | ThermoFisher | 61965059 | Digestion buffer |
| HEPES (1M) | ThermoFisher | 15630056 | Digestion buffer |
| Bovine serum albumin | Sigma | A4503 | Staining buffer |
| DAPI | ThermoFisher | D1306 | Histology and FACS |
| Dow Corning high-vacuum s. grease | Sigma | Z273554 | Sample mounting |
| Menzel cover slips (24x32 mm, #1) | HUBERLAB | 10.0360.22 | Histology |
| SuperFrost Microscope Slides | ThermoFisher | 12134682 | Histology |
| 16% Paraformaldehyde | EMS | 15710 | Tissue fixation |
| RNeasy Plus Micro Kit | Qiagen | 74034 | RNA extraction |
| 2-Mercaptoethanol | Sigma | M3148 | RNA extraction |
| Nonstick, RNase-free Microfuge Tubes | ThermoFisher | AM12450 | RNA extraction |
| Falcon Cell Strainers | ThermoFisher | 08-771-2 | Flow cytometry |
